# Different dispersal rates and declining climate suitability shape future vegetation compositions across the Arctic

**DOI:** 10.1101/2025.08.22.671233

**Authors:** Ronja Schwenkler, Ulrike Herzschuh, Luca Zsofia Farkas, Boris Schröder, Simeon Lisovski

## Abstract

**Aim:** We investigate how species-specific dispersal abilities might influence future Arctic plant distributions and large-scale dynamics at the boreal forest – tundra boundary until 2100

**Location:** circumpolar terrestrial Arctic (boreal forest, taiga and tundra)

**Taxon:** 1,550 plant species

**Methods:** We developed climate-driven species distribution models (SDM) to predict species-specific emerging climate niches under different climate scenarios. The model was parameterized using occurrence data from the Global Biodiversity Information Facility database (GBIF) and temperature and bioclimatic variables from the CHELSA data set. Dispersal rates were assigned to each species using a trait-based approach and were used to predict future habitat with a distance-based probability over time.

**Results:** Plant species are predicted to occupy on average only 12.3% (1.5 – 53.9 95% CI) of their emerging climate niches, with half of the species unable to colonize new habitat by 2100 due to limited dispersal distances. In dispersal limited predictions migration to higher altitudes played a greater role than northward shifts. Decolonization by species (extirpation) due to decreasing climate suitability had a larger effect on species composition change compared to dispersal limitations. Boreal tree species were predicted to expand into the tundra shrinking the treeless areas.

**Main conclusions:** Future plant species distributions and resulting large-scale compositions are affected by species-specific dispersal rates. Even though new suitable niches emerge prominently towards the north, higher altitudes might be more relevant given their accessibility by dispersal over the next century. Although climate niche dynamics could support higher plant species richness across the Arctic, overall richness is expected to decline with climate warming due to dispersal limitations. The colonization of new habitats via dispersal in combination with the decolonization of former habitats due to declining climatic suitability on species level are predicted to cause large-scale changes in species composition, especially at the boundary between the boreal forest and the tundra biome.

## 1. Introduction

Over the last decades, the temperature in the Arctic has increased two to four times faster compared to the global average, and the speed of warming might even accelerate in the near future (Rantanen et al. 2022). As a result, species’ ranges and pehnologies are changing across the entire Arctic (Parmesan et al. 2006). The northward and upward migration of plant species is a frequently reported response to the changes in the climate niche (Iverson and McKenzie 2013; Stewart et al. 2018; Niskanen et al. 2019; Roland et al. 2019). The advance of the boreal forest into the tundra biome is of particular interest, since it might threaten tundra biodiversity and function (Hofgaard et al. 2012) and might lead to a large-scale biome shift (Beck et al. 2011; Berner and Goetz 2022; Montesano et al. 2024). And although forest expansion and shrubification are expected to cause significant changes to the ecosystem (Mod and Luoto 2016), it remains uncertain how Arctic plant communities as a whole will shift their distribution in the near future, and how these changes might drive large-scale structural transformations across the Arctic.

The boreal forest, also known as the taiga, and the tundra represent two of the most extensive and distinct biomes in the Northern Hemisphere, each characterized by unique biodiversity and ecosystem functions. The boreal forests are dominated by coniferous trees, with spruces, pines and firs in northern North America (Nearctic) and mostly larches in eastern Eurasia (Palearctic) (Herzschuh 2020). In contrast, the treeless tundra is mostly covered by mosses and lichens, dwarf shrubs, and forbs (Zhang et al. 2013), being shaped by extreme cold, permafrost, and short growing season (Shaver 1995). Shrubification of the tundra would reduce the albedo of the landscape, thereby contributing to warming and release of the large amounts of carbon stored in the permafrost soils (Myers-Smith et al. 2011; Bonfils et al. 2012; Loranty and Goetz 2012; Pearson et al. 2013; Plaza et al. 2019). During the last decades an obvious change within the biomes has been driven by the longer growing season with increased plant productivity, leading to a greener Arctic (Phoenix and Bjerke 2016; Lara et al. 2018; Arndt et al. 2019; Tømmervik and Forbes 2020). More recently, this has locally been reversed by frequent summer droughts and wildfires, decreasing plant productivity (Phoenix and Bjerke 2016; Tømmervik and Forbes 2020). For the future, we expect that changes in species composition due to range shifts, both within and across the biomes, will have even stronger effects on the ecosystems, especially the tundra. Today, large parts of the tundra are already suitable for growth of shrubs and trees, and their potential distribution is predicted to cover almost the entire tundra by the end of the century (Zhang et al. 2013). While boreal forests expand at their northern edge, they shrink at their southern edge, being replaced by other biomes, which leads to a biome shift (Beck et al. 2011). However, treeline migration seems relatively slow, significantly lagging behind the climate niche (dispersal lag) (Primack and Miao 1992; Svenning and Sandel 2013; Kruse und Herzschuh 2022), e.g. with thermal velocity exceeding species’ dispersal by 2 – 4 km/year according to a past simulation (Zani et al. 2023). After the last glacial maximum, the average dispersal rate of 140 studied European plant species was 1.6 km/year, considerably varying with dispersal mode, as e.g. brown bears (*Ursus arctos*) can transport seeds about 6 km/year (Cunze et al. 2013). Since the last glaciation, the North American jack pine (*Pinus banksiana*) and black spruce (*Picea mariana*) have migrated 19 km and 25 km per century, respectively (Payette et al. 2022). During the 19^th^ and 20^th^ century, the White spruce *(Picea glauca)* treeline moved up to 100 m into the tundra in Alaska (Suarez et al. 1999). For the 21^st^ century, recent models forecast a much faster advance of the treeline, e.g. the individual-based model developed by Kruse und Herzschuh (2022) predicts a migration of larches of up to 3 km/year under climate warming.

The ability of a plant species to colonize habitats within its climatically suitable niche depends on various factors (McNichol and Russo 2023), but it is primarily constrained by its dispersal capacity (Eriksson 2000; Kruse et al. 2019). Plants can be dispersed by wind (anemochory), water (hydrochory) and animals (zoochory), or the seeds simply fall to the ground, thus the dispersal distance is only about the height of the parent’s stem (local non-specific) (Bierzychudek 1982; Hughes et al. 1994; Lososová et al. 2023). Besides these dispersal modes, other plant traits, e.g. the height and growth form, determine the possible dispersal distance of a species (Lososová et al. 2023). Herbs occurring in deciduous forests often do not have special dispersal mechanisms, thus their seeds only disperse locally, and vegetative reproduction is common (Bierzychudek 1982). Short and cold summers hinders sexual reproduction (Bell and Bliss 1980), thus tundra plants reproduce only infrequently at their northern distributional limit (Molau 1993). Still, most species reproduce via seeds, but can additionally reproduce vegetatively (Billings 1987) or via vivipary (Molau 1993), resulting in very local dispersal as the next generation evolves close to the parent plant. Considering dispersal constraints in predictions of future climate niches has the potential to provide more realistic scenarios of potential changes in distributions of single species and entire communities (Jaeschke et al. 2013).

Species distribution models (SDMs) combine species occurrence data with environmental variables to predict distributions across landscapes, also extrapolating in space and time (Elith and Leathwick 2009). SDMs are based on the assumption that species occurrences are in equilibrium with their climate niche, however this assumption is usually violated when species shift their range under changing climate conditions (Václavík and Meentemeyer 2011). Beside the dispersal lag, which refers to limited dispersal abilities, there is the establishment lag, which implies that the species are facing difficulties to establish due to e.g. competition with the present community, and the extirpation lag, describing the phenomenon that species will stay in their old apparently unsuitable habitats for a longer time than expected as living individuals do not die instantly and seeds are still stored in the soil (Alexander et al. 2018). From these different time lags, a disequilibrium between vegetation and climate results, which is difficult to predict (Svenning and Sandel 2013) but is still important to be included in models forecasting spatio-temporal changes (Alexander et al. 2018). Nevertheless, most species distribution models do not account for these lags. Implementation of these constraining mechanisms is possible in models that are based on cellular automata(Engler et al. 2012; Zani et al. 2023), but difficult to tackle in statistical (correlative) models. Probably as pioneers, Shipley et al. (2022) implemented dispersal in a statistical SDM, calculating the probability of dispersal based on supplied dispersal rates using exponential distributions, while Jaeschke et al. (2013) had cut their predictions of Odonata range shifts using ArcGIS.

In this study, we applied statistical SDMs implementing species-specific dispersal rates based on plant trait information, and conducted an additional analysis regarding the extirpation lag. Our goal was to investigate how the different dispersal abilities of species might impact the large-scale dynamics at the boreal forest – tundra boundary until 2100. We hypothesized that constraining dispersal rates lead to smaller distribution changes compared to the potential of emerging habitats. However, even though there are relatively small changes within species, the consequent changes in species composition may still cause structural changes across the Arctic. Considering a large-scale approach, we separately modeled the distribution of 1,550 terrestrial plant species, to predict (1) emerging suitable climate niches across the Arctic, (2) the area that can potentially be colonized by plant species given the dispersal constraints, and (3) spatial patterns of range shifts with a focus on northward movements. By combining the future prediction of all species ranges, we test the hypothesis that (4) changes on species level can lead to large-scale changes at the boundary between the boreal forest and the tundra biome until 2100.

## 2. Material and Methods

### 2.1 General modeling approach

We applied climate-driven species distribution models (SDM) to predict emerging niches for plant species across the Arctic boreal forest, taiga and tundra. Based on trait specific dispersal distances, we compared all emerging areas with the areas that could effectively be reached and potentially be colonized given species-specific dispersal capacities. Covering the entire terrestrial Arctic with a 25 km x 25 km resolution, our model predictions aim to generate insights into landscape-scale dynamics of species distributions, how large-scale patterns may change due to ongoing climate change, and how species-specific dispersal affects predictions. All analyses were performed using R Statistical Software (version 4.3.1; R Core Team 2023), the model code is available and a release will be published on Zenodo after acceptance). For more details see the ODMAP protocol (Appendix S1) according to the criteria provided by Zurell et al. (2020) and further information in the Appendix S2, S3 and S4 in Supporting Information.

### 2.2 Occurrence data, background data and environmental variables

To define the current climate niches of Arctic plant species (circumpolar Arctic, delimited by boreal forest/taiga and tundra polygons; Olson et al., 2001), we retrieved all available presence records from the Global Biodiversity Information Facility (GBIF) using the package rgbif (v.3.7.8; GBIF Occurrence Download, 26 March 2024). Occurrence data were cleaned following GBIF protocols (GBIF Secretariat, 2021). We removed records with missing or erroneous coordinates or species names, fossil records, and museum specimens. Only records collected after 1970 and with a coordinate uncertainty < 5 km were retained. The final dataset is registered with GBIF as a derived dataset (doi:10.15468/dd.y4uq5q) and archived on Zenodo (https://zenodo.org/records/16925866).

To reduce sampling bias, we first applied grid-based thinning by retaining one occurrence per 5 × 5 km environmental grid cell. Because records are strongly clustered, with a particular bias towards Europe (Figure 1a), we additionally applied spatial filtering (Veloz, 2009; Kramer-Schadt et al., 2013): random sample points were generated with an approximate 150 km interval, and the closest occurrence point was retained for each (see Appendix S2, cf. Figure S2.3 for a comparison with models without this step). The filtered dataset was randomly split into training (75%) and test data (25%). Finally, we retained only species with ≥150 occurrences for training, after the described thinning and filtering processes. Across species, the median number of training records was 680 (95% CI: 153–1777) and of test records 227 (95% CI: 51–592). Species-specific details are provided in Appendices S3 and S5.

**Figure 1:**
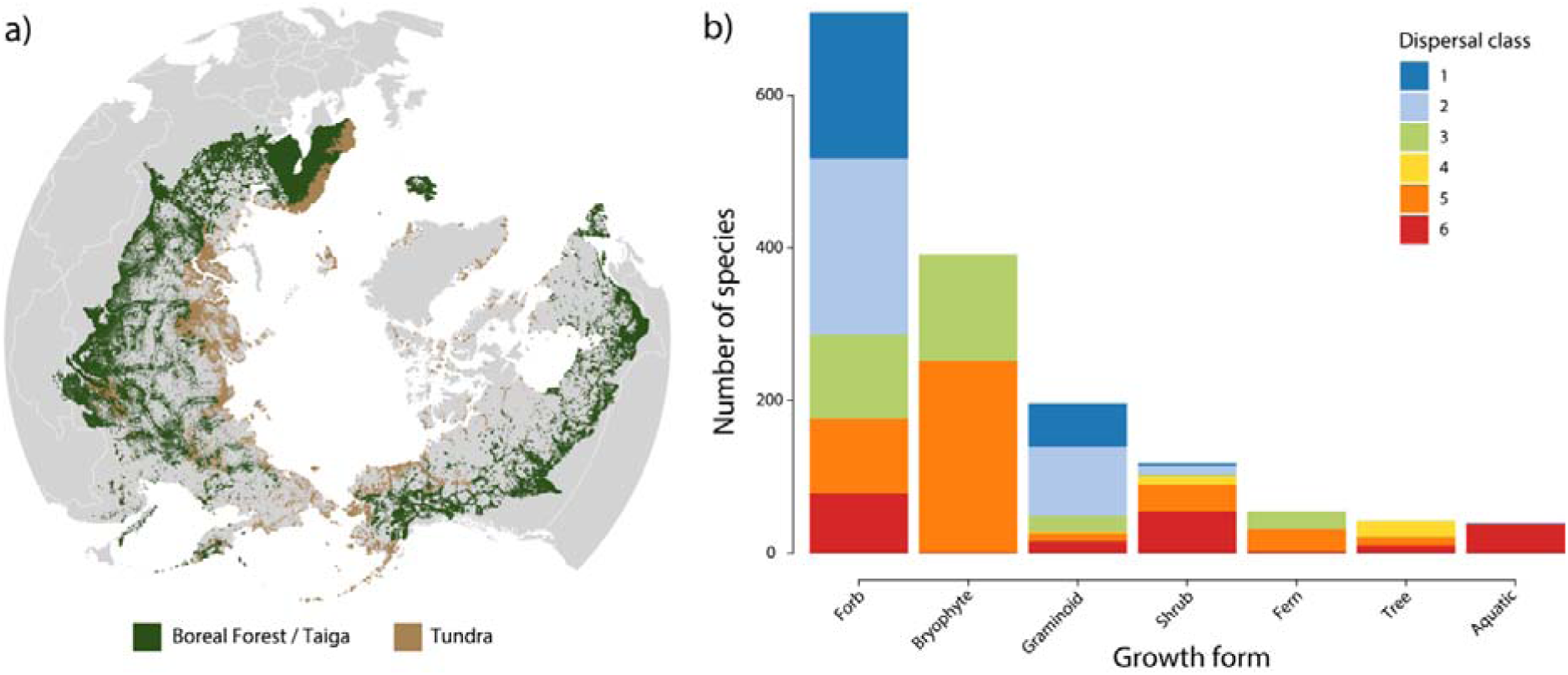
(a) Spatially thinned Plantae occurrence records (n = 4,445,063) from GBIF (Global Biodiversity Information Facility) for the boreal forest/taiga and tundra (defined following Olson et al. 2001). (b) Number of Plantea species from the thinned GBIF records across six different dispersal classes and growth forms. The dispersal distance classes were assigned according to Lososová et al. (2023) and describe the maximum dispersal distance of 99% of the seeds ranging from 1 m (class 1) to 1,500 m (class 6). Map created with Natural Earth. [double column]

Background data were generated following Senay et al. (2013). We defined the geographical extent by applying a buffer around presence points and then profiled the background using a one-class support vector machine classifier. Across species, the median number of background points was 18,606 for training (95% CI: 18,389–18,717) and 6,202 for testing (95% CI: 6,130–6,239). Species-specific numbers of training and test points are reported in Appendix S3 and S5.

As predictor variables, we used the CHELSA v2.1 climate dataset (Karger et al., 2017; 2021) at 30 arc-second resolution, resampled to 5 × 5 km in an equal-area projection (downloaded March 27, 2024). For model calibration, we used historical monthly climate data (1981–2010). For future projections, we used outputs from the global climate model GFDL-ESM4 (CMIP6). This model was selected based on the CHELSA technical guidelines (CHELSA 2021), which follow the prioritization scheme of the Intersectoral Impact Model Intercomparison Project (ISIMIP 2021). The ISIMIP is an international network of climate-impact modelers that provides a framework for consistently projecting the impacts of climate change across affected sectors and spatial scales (Schellnhuber et al. 2013). Predictor variables included maximum and minimum air temperature as well as bioclimatic indices (Hijmans et al., 2005). The bioclimatic variables represent seasonal and annual statistics of temperature and precipitation (Hijmans et al. 2005). We retained bio1 (annual mean temperature) and bio12 (annual precipitation) for all species, and added further predictors after collinearity filtering. Variable importance across models is shown in Appendix S2 (Figure S2.1), and species-specific predictors are listed in Appendix S3.

Climate data were provided as monthly means for 30-year periods and aggregated to annual means. We used the periods 2011–2040, 2041–2070, and 2071–2100 (hereafter referred to as 2040, 2070, and 2100) under three emission scenarios: ssp126, ssp370, and ssp585. These scenarios combine Representative Concentration Pathways (RCPs) describing radiative forcing levels of 2.6, 7.0, and 8.5 W/m² (IPCC, 2014) with Shared Socioeconomic Pathways (SSPs; O’Neill et al., 2017). ssp1 represents a sustainable future, ssp3 regional rivalry and fragmentation, and ssp5 fossil-fueled development with population growth in high-income countries and decline in developing countries.

### 2.3 Predicting future niche suitability with species distribution models

To predict future niche suitabilities, we used the MaxEnt model (Phillips et al. 2006), implemented in R package ‘dismo’ (v. 1.3.14). MaxEnt is considered to have high predictive accuracy (Merow et al. 2013) even with presence-only data, using background data instead of (pseudo-)absences (Phillips et al. 2006). We predict the climate suitability (0 – 1) of the present time slice (1981 – 2010) for all 1,550 plant species based on the thinned occurrence points, the generated background data and the climate data (for settings see Appendix S2). To transfer the continuous output (0 – 1) into binary values (0 or 1) we used the MaxEnt prevalence threshold “10 percentile training presence” which describes that 90% of all input occurrence points are included in the probabilities above the threshold (for values see Appendix S3).

### 2.4 Implementation of dispersal constraint

Dispersal was implemented by considering species-specific dispersal rates. For each species, the traits growth form, height, habitat and dispersal-related diaspore features were collected from various databases, literature and websites (Farkas et al. (2025), and Appendix S4). Based on these traits, the dispersal mode (e.g. anemochory, zoochory) and the resulting dispersal distance class (1 – 6) were assigned to each species according to Lososová et al. (2023) (Figure 1b). The dispersal distance classes describe the maximum dispersal distance of 99% of the seeds ranging from 1 m (class 1) to 1,500 m (class 6). For each future scenario, based on distance to the closest occurrence (always in relation to the binary map of the presence time slice) and a sigmoidal probability function parameterized with the 50% and 99% dispersal distances, the MaxEnt suitability for each cell was set to zero if a random number between 1 and 0 was above the sigmoidal probability value (and original value was kept for random numbers below). For comparison, we also ran a scenario without the implementation of the species-specific dispersal rates. For this “unconstrained” scenario, we used a normal distribution probability function with mean = 1,500 km and sd = 250 km.

### 2.5 Analyzing the shifts in boreal forest–tundra distribution and community composition

To study large-scale dynamics of boreal forest and tundra, we first identified tree species that do not occur as shrubs or dwarf forms (Appendix S10). For each grid cell, we calculated the ratio of these boreal indicator tree species to the total number of species. This ratio was then converted to a binary classification: “no trees” (>0% trees absent) representing tundra, and “trees present” (>0% trees) representing boreal forest/taiga. Cells classified as boreal forest were further assigned to the Nearctic or Palearctic based on WWF Terrestrial Ecoregions (Olson et al., 2001). Agreement with established ecoregions was high: 89% for the Nearctic boreal forest/taiga and 93% for the Palearctic. The “no tree” areas, however, covered only 63% of the tundra ecoregion. This mismatch arises because our classification assigns a cell to boreal forest if even one typical tree species is predicted present, whereas tundra in reality is defined by characteristic species assemblages. In the following, we use “tundra” for our “no tree” area and “boreal forest” for the biome “boreal forest/taiga,” while noting that correspondence with true ecoregions is not complete.

To analyze changes in species composition, we stacked all species distribution maps and calculated the Jaccard dissimilarity index, comparing projections for 2040, 2070, and 2100 to the baseline of 2010. Values range from 0 (total similarity) to 1 (total dissimilarity). We also partitioned beta diversity into turnover (species replacement) and nestedness (species loss or gain without replacement) using the betapart package (v.1.6.1; Baselga et al., 2025). Finally, we included a “no decolonization” scenario, where species may colonize new cells but cannot disappear from cells that become climatically unsuitable (extirpation).

### 2.6 Credible intervals and significance testing

Slopes were estimated with linear regression using the function ‘lm’ from ‘stats’. We then simulated 1000 values from the joint posterior distribution of the parameters of our linear model using the function ‘sim’ from the package ‘arm’ (v.1.14.4) for deriving the 95% credible intervals of the slope. Significant differences were tested with the Wilcoxon test using the function ‘wilcox.test’ from the package ‘stats’ (v. 4.3.1).

## 3. Results

### 3.1 Overall SDM performance

Averaged over the SDMs for all 1,550 species the AUC was 0.89 ± 0.07 for training data and 0.87 ± 0.09 for test data. The Boyce Index (ranging from –1 to 1) was 0.81 ± 0.15 for training data and 0.69 ± 0.26 for test data. For SDM calibration maps (1980-2100) along with GBIF occurrence points see Appendix S5.

### 3.2 Dispersal-limited colonization of new habitats

The area of new suitable habitat increased over time and under more extreme climate scenarios (Figure 2a). By 2100, under scenario ssp585, species were predicted to gain a median of 4 × 10D km² of new suitable habitat within the study region (24 × 10D km² in total). Similar gains were projected under ssp126 (3.9 × 10D km²) and ssp370 (4.1 × 10D km²). However, 844 species (54%) were not able to colonize any of these new areas. Across all scenarios and time steps, the remaining species could reach only 12.3% (95% CI: 1.5–53.9 %) of their potential new habitats through dispersal (ssp126: 9.3–18.1%; ssp370: 9.6–14.9%; ssp585: 9.4–14.3 %).

**Figure 2:**
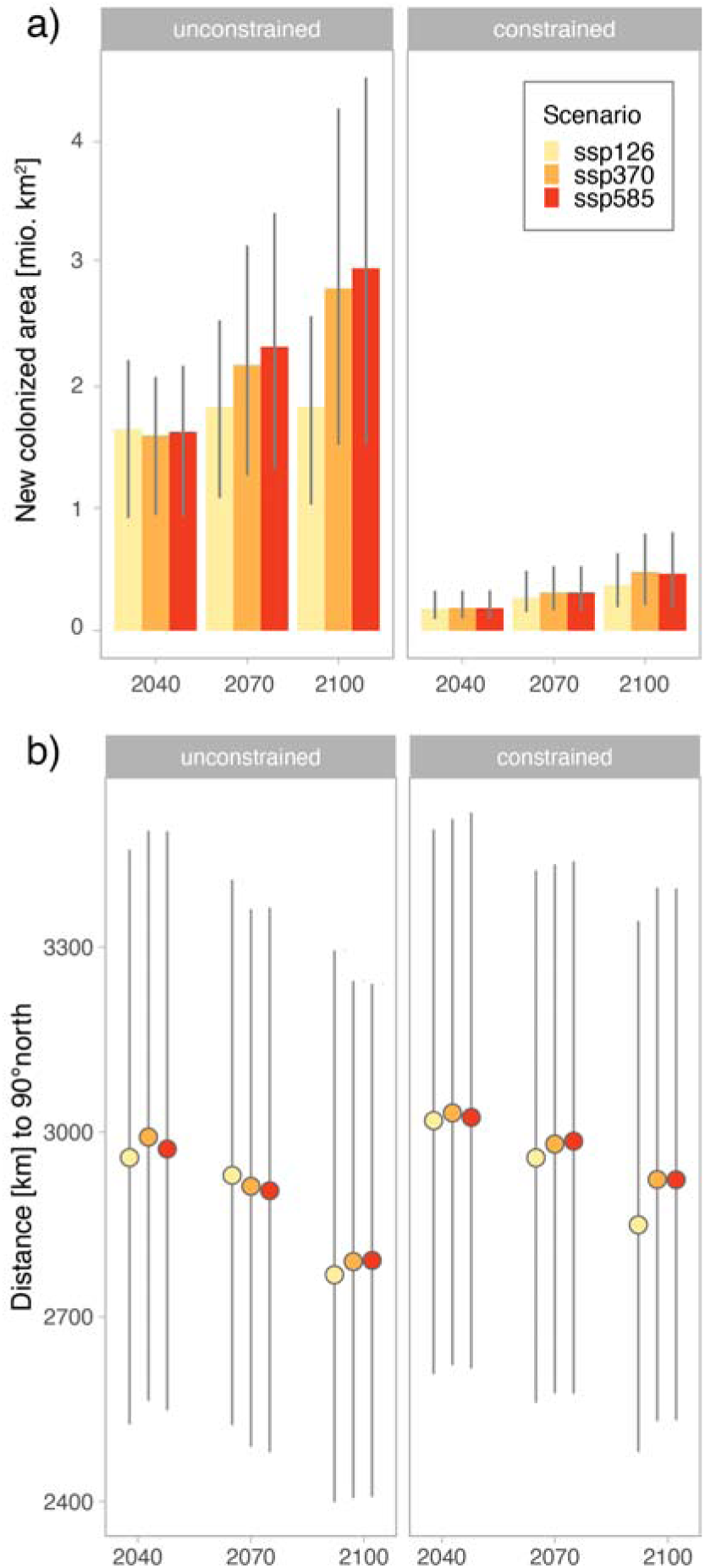
(a) New colonized area [km2] for the unconstrained and species-specific dispersal constrained scenario (compared to 2010) averaged across the 1550 arctic plant species. (b) Distance [km] to 90° north of new suitable cells (unconstrained) and newly colonized cells (dispersal constrained) (delta to time slice before) averaged over all 1550 species. New cells refer to areas which indicated “absence” in 2010 and “presence” in the respective future time slice. Circles with error-bars show the median and the 25% and 75% quantiles. [single column]

Dispersal ability strongly constrained colonization. Species were assigned to six dispersal distance classes based on traits, with 57% limited to <150 m/year. Colonization ranged from 0% in the lowest class to 36% (mean) in the highest (class 6; Figure 3a). Because medians were often zero, means are reported here. Dispersal mode was the main determinant of class assignment (Lososová et al., 2023). Species with very short dispersal distances (1–5 m, class 1) were unable to colonize new habitats. Myrmecochorous species (class 3) could colonize only 2.5 × 10⁻D% of their potential area, whereas anemochorous species reached 8.7%. Zoochorous and hydrochorous species (up to 1500 m/year) colonized much larger shares, 33.2% and 39.3%, respectively (Figure 3b).

**Figure 3:**
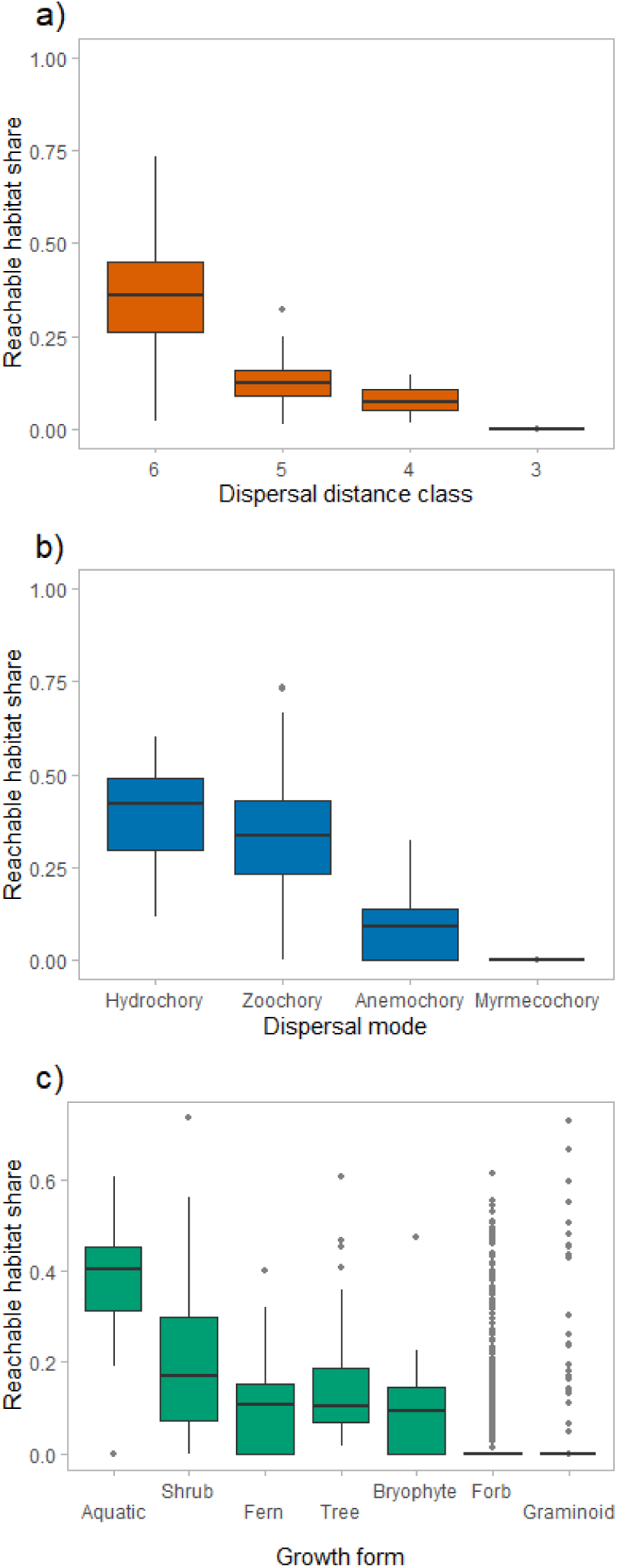
Reachable habitat share, i.e. new colonized cells in 2100 (climate scenario ssp585) compared to 2010 for the dispersal constrained scenario divided by the new colonized cells in 2100 (climate scenario ssp585) compared to 2010 for the unconstrained scenario for the (a) dispersal distance class, (b) dispersal mode and (c) growth form. [single column]

Growth form also influenced colonization potential (Figure 3c). Aquatic species, mostly hydrochorous (class 6), colonized the largest share of new habitats (37.6%). Among terrestrial plants, shrubs (19.7%) and trees (15.9%) performed best, followed by ferns (9.4%) and bryophytes (8.5%). Forbs (5.4%) and graminoids (4.3%) had the lowest colonization success. Thus, boreal forest growth forms generally had greater colonization potential than tundra species. Results for ssp126 and ssp370 are shown in Appendix S12, Figure 4).

**Figure 4:**
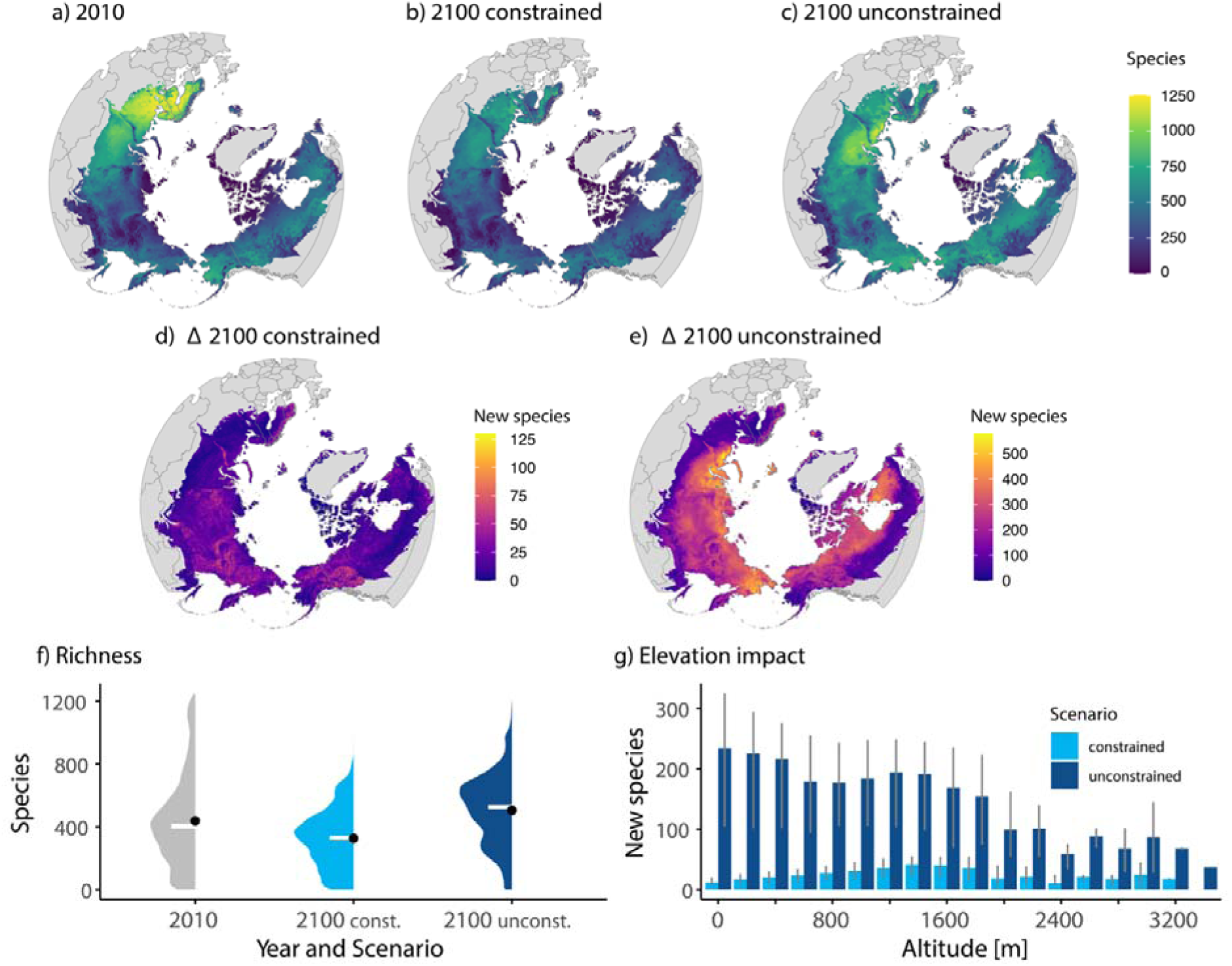
(a) Number of predicted (ssp585) plant species for present (2010), with (b) dispersal constraint and (c) unconstratin future (2100) distributions. Number of only new colonizing species by 2100 compared to 2010, for (d) dispersal constrained and (e) unconstrained scenario. (f) Violin plot with median of number of all species per cell (data from a-c). (g) Number of new species over different altitudes for the constrained and unconstrained predictions. Error bars show the 25% and 75% quantiles. [double column]

### 3.3 Distribution of new habitats

Our analysis revealed distinct spatial patterns in the emergence of new habitats under a warming climate. The model predicted a northward shift of emerging habitats over time for the unconstrained scenario (ssp126: slope (absolute values) 1.76 km/year, 1.73 – 1.79 95% CI; ssp370: 3.18, 3.16–3.19 95% CI; ssp585: 3.03, 3.01–3.04 95% CI) (Figure 2b). The habitats that could be reached under dispersal limitation showed a northward shift as well (ssp126: 2.16 km/year, 2.10–2.22 95% CI; ssp370: 1.53, 1.48–1.58 95% CI; ssp585 1.49, 1.44–1.54 95% CI). The climate scenarios did not exhibit a linear impact on the shift.

Species richness showed a general decrease towards the north, with a species hotspot in Scandinavia and the lowest richness in Greenland (Figure 4a). The maximum cell species richness was predicted to decline from 1251 in 2010 to 1006 (constrained) and 1202 (unconstrained) by 2100 (Figure 4b, c). The mean number of species per cell was projected to decrease from the current 437 (median 403) to 326 (median 328) in the constrained scenario and increase to 505 (median 526) in the unconstrained scenario by 2100. (Figure 4f).

Colonization patterns were strongly influenced by dispersal abilities. The maximum number of new colonizing species per cell was 130 in the constrained scenario and 578 in the unconstrained scenario (Figure 4d, e). Especially for the unconstrained scenario, most colonizations occurred in the northern part of the Arctic. Altitude also played a role in colonization patterns, with most colonizations occurring around 1200 m in the constrained scenario and around 0 and 1400 m in the unconstrained scenario (Figure 4g). For scenarios ssp126 and ssp370, see Appendix S12, Figure 4.

### 3.4 Shifts in boreal forest–tundra distribution and community composition

In the constrained scenario, the share of boreal indicator tree species changed little over time (Figure 5a). In contrast, the “no tree” area declined from 27% to 12% in the unconstrained scenario, accompanied by an increasing share of boreal indicator trees (Figure 5b). Dispersal dynamics also shifted: while slow dispersers (classes 1–3) dominated 78% of cells in 2010, by 2100 more than half of the species were fast dispersers (classes 4–6) in 66% of cells in the constrained scenario (Figure 5c). The Palearctic boreal forest colonized more new cells than the Nearctic, whereas the Nearctic showed a significantly stronger northward shift (Wilcoxon test: p < 2.2e-16; Figure 5d). All these changes were more pronounced in the unconstrained than in the constrained scenario.

**Figure 5:**
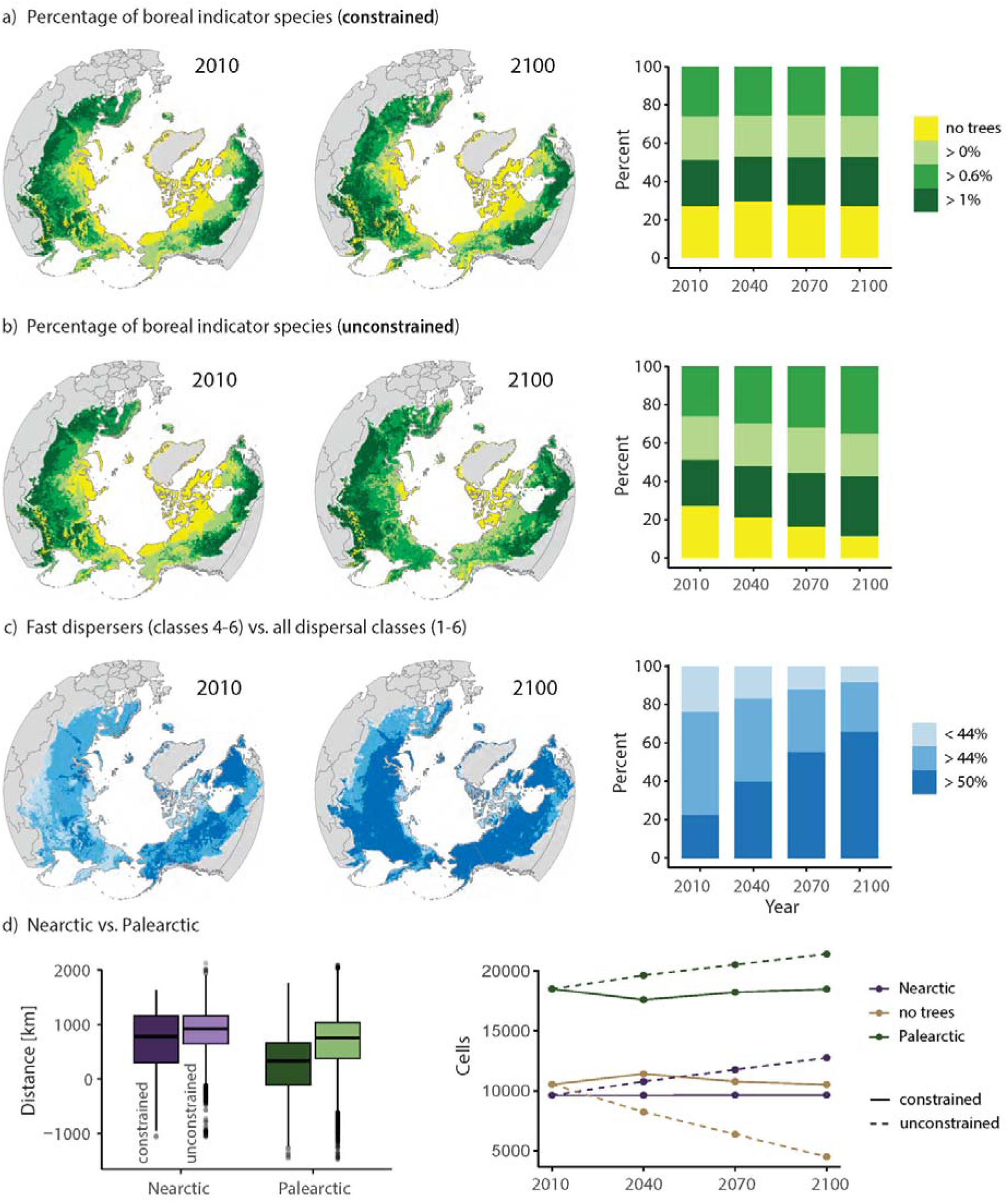
Temporal dynamic of the boreal forest – tundra boundary over time (2010 to 2100) for climate scenario ssp585, visualized as the percentage of boreal indicator tree species in total number of plant species per cell for (a) dispersal constrained and (b) unconstrained predictions. c) Percentage of quick dispersing species (dispersal distance classes 4-6) among all species (classes 1-6). (d, left) Northwards shift of the boreal forest communities across the Nearctic and the Palearctic for constrained and unconstrained predictions. (d, right) Number of cells occupied by Nearctic and Palearctic boreal forest communities and treeless communities over time for dispersal constrained (solid line) and unconstrained predictions (dashed line). Assignment to Nearctic, Palearctic and treeless communities were made according to their geographic position in the realms and the similarity to the biomes of the boreal forests / taiga and the Arctic tundra (Olson et al. 2001). [double column]

Species composition diverged increasingly from 2010 over time (Figure 6). By 2100, median dissimilarity reached 0.29 (95% CI: 0.09–0.78) in the constrained scenario and 0.53 (0.35–0.92) in the unconstrained scenario. To separate the effects of colonization and decolonization, we added a “no decolonization” scenario. Here, dissimilarity remained low (median 0.05, 95% CI: 0.003–0.29), highlighting that climate-driven extirpations were a major driver of compositional change. Indeed, the unconstrained scenario produced on average 224% of the constrained dissimilarity, while the no decolonization scenario produced only 26%.

**Figure 6:**
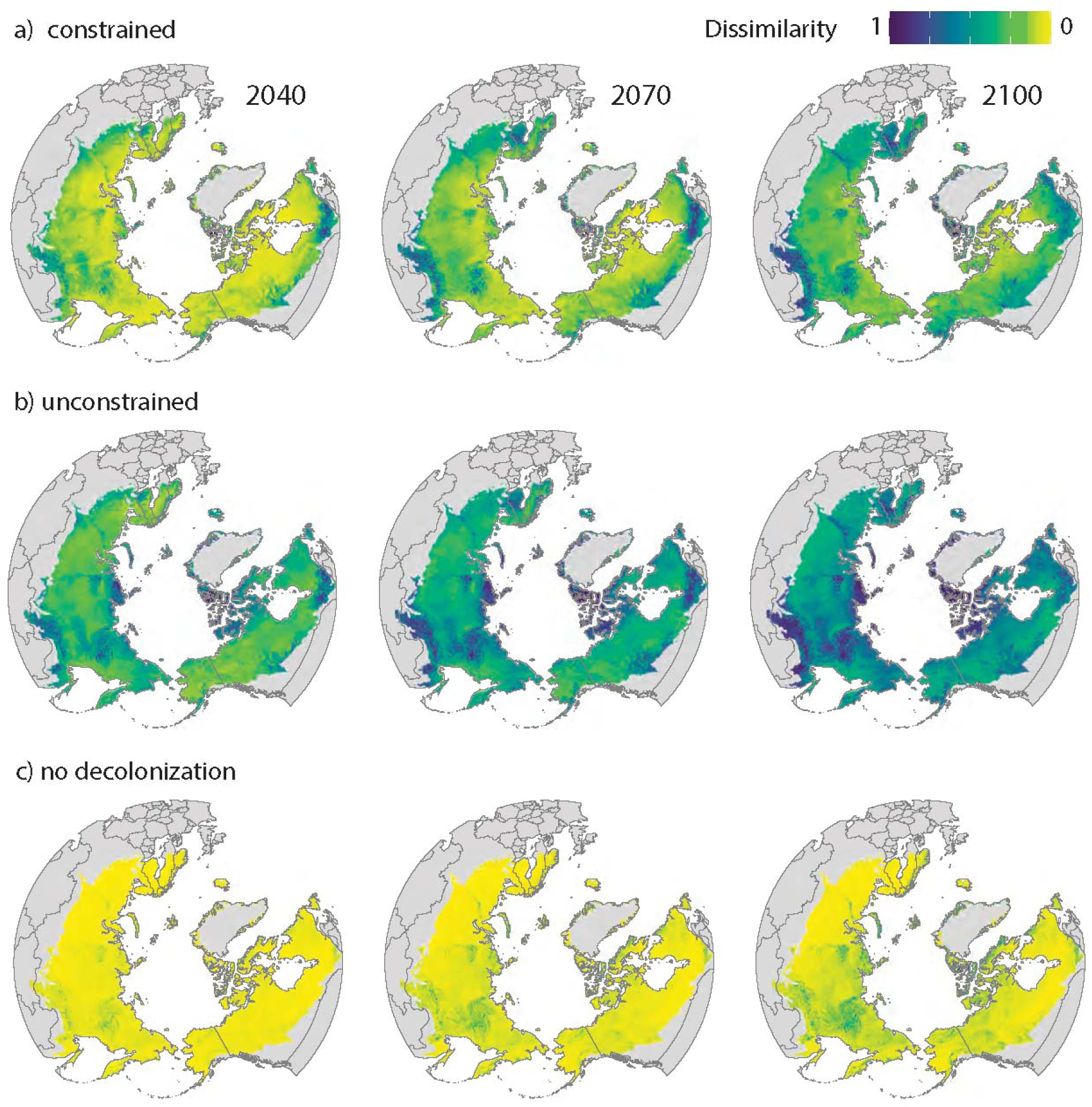
Effect of dispersal constraint and extirpation assumptions on predicted plant communities: Dissimilarity (Jaccard index from 0 o 1) of species composition compared to 2010 over time for (a) constrained, (b) unconstrained, and (c) no decolonization (and constraint) predictions. In the no decolonization predictions species may only colonize new cells but may not disappear if the climate niche becomes unsuitable. [double column]

Partitioning beta diversity further showed that nestedness (species loss or gain without replacement) dominated under dispersal-constrained conditions, largely due to losses in the south (Appendix S11, Figure S11.1). Under unconstrained conditions, turnover (species replacement) became equally important, as colonizations could compensate losses. In the no decolonization scenario, changes were exclusively due to colonization, and thus entirely nestedness.

## 4. Discussion

We examined how dispersal ability influences future distributions, community composition, and the boreal forest–tundra boundary by projecting emerging climate niches under different scenarios until 2100. Our results support the hypothesis that dispersal strongly constrains range shifts. Only a fraction of emerging habitats is reachable, yet this may still drive substantial shifts of the forest– tundra boundary.

### 4.1 Limitations and validation

Our study focuses on dispersal constraints in climate-driven range shifts, but other processes can also shape outcomes. Biotic interactions influence both the rate and direction of range shifts (HilleRisLambers et al., 2013), competition may slow down expansion at lower range limits, while facilitation may accelerate upward shifts (Ettinger & HilleRisLambers, 2017). Competitive tundra vegetation can restrict shrub expansion (Angers-Blondin et al., 2018; Armitage & Jones, 2020), while establishment lags caused by e.g. competition can delay colonization (Alexander et al., 2018). Tall deciduous shrubs, in contrast, often expand due to strong competitive abilities (Mekonnen et al., 2021; Usinowicz & Levine, 2021). Extirpation lags (“extinction debt”) may also delay species loss, supported by long-lived individuals or soil seed banks (Tilman et al., 1994; Alexander et al., 2018).

Comparing our results with a no-decolonization (extirpation) assumption (Figure 6c) led to less pronounced changes, clarifying that decolonizations due to declining climatic suitability can substantially impact species composition, and hence extirpation lag can have an effect. However, because empirical data for these processes are scarce, we only included dispersal lag explicitly and focused most analyses on colonization. For species’ range shifts we considered both colonizations and decolonizations, but ignored establishment and extirpation lags. This assumption was partly justified by observations of equal leading– and trailing-edge shifts (Rumpf et al., 2019a), though responses vary among species (Rumpf et al., 2019b; Robeck et al., 2024). Our calibration, based only on occurrences within the Arctic, may have missed suitable climate analogues outside the study area. Nevertheless, comparison with a “no decolonization” scenario (Figure 6c) confirmed that extirpations due to declining climate suitability strongly affect community composition.

Our calibration of species’ climate niches relied solely on occurrence records from within the Arctic study area (boreal forest and tundra). As a result, the modeled suitability represents only this, however often large, portion of each species’ niche. Potential suitability inferred from climate analogues outside the study area could not be captured. Consequently, some climatically suitable sites may have gone undetected. Furthermore, southern species with range shifts into the study area were ignored.

Despite the coarse resolution and exclusion of non-climatic factors such as soil fertility (Schibalski et al., 2014), our models reproduced known patterns. Assigned dispersal classes (Lososová et al., 2023) were in line with literature estimates (Appendix S6). Current distributions matched reported ranges (e.g., Picea glauca in Alaska; Ohse et al., 2009; Appendix S7). Future projections aligned with observed northward and upward shifts of thermophilic species such as *Alnus glutinosa* in Scandinavia (Kullman, 2008). The modeled richness gradient—with a hotspot in Scandinavia and lowest richness in Greenland—also reflects published patterns (Mutke & Barthlott, 2005; Pironon et al., 2020). Field validation in Ulukhaktok (Northwest Territories, Canada) further supported our results. Of 49 recorded species, 36 occurred in our dataset, and 32 (86%) were correctly predicted by the model for 2010–2040 despite sparse GBIF coverage in this region (Appendix S8).

### 4.2 Multiple species-specific responses to warming

Species range shifts to the North were not predicted as pronounced as expected. We found a northward shift in the predicted emerging habitats (as reported by Stewart et al. 2018; Roland et al. 2019), but considering different dispersal abilities, only a small share or even nothing of the predicted new habitat area could be reached. Consequently, dispersal abilities significantly impact the future of individual species by defining their capacity to follow their climate niche under changing climate conditions. However, even trees, which belong to the faster dispersers, will probably not be able to keep up with the predicted shift of their climate niche until 2100 (Hanberry 2024).

Besides the northward shift, another response of species to increasing temperatures is the colonization of higher altitudes (MacDougall et al. 2021). In our model prediction, we also observed that under dispersal limitation the peak of colonizations occurred at higher altitudes than under unconstrained conditions, where a colonization peak occurred at 0 m. This suggests that suitable climate niches at higher altitudes may serve as more relevant refugia due to higher accessibility, while those at higher latitudes remain largely unreachable. In response to warming, particularly non-boreal species migrate to higher elevations, thereby also responding to boreal taxa that tend to expand their distribution ranges (MacDougall et al. 2021).

Dispersal abilities could substantially determine to which extent richness decline can be counterbalanced, as species richness was even predicted to increase on average without dispersal limitation while it decreased when considering dispersal. In reality, due to the effect of competition, intermediate dispersal distances allow for highest richness, as small dispersal distances only enable few colonizations while large dispersal distances lead to homogenization of the metacommunity and the dominance of the most competitive species (Mouquet and Loreau 2003).

The compositional changes we observed were driven not only by northward migration but even more by local disappearances, as suitable habitats declined in southern areas under climate change. This highlights the importance of considering extirpation lag in improving model predictions. In reality, southern declines of tundra specialists may be further reinforced by low competitive ability, as these species risk being outcompeted by generalists expanding northward (Callaghan et al., 2004; 2005; Pykälä, 2017).

### 4.3 Shifts in community composition

Species-level variation in climate responses is reflected at broader scales. In our study, boreal forest area expanded while tundra contracted, as many tundra cells in 2010 shifted to boreal forest by 2100. This shift is driven mainly by fast-dispersing boreal species that migrate northwards and colonize treeless areas. At the same time, species decolonized southern habitats where climate suitability declined, altering the composition that defines both biomes. Because tundra species generally disperse more slowly than boreal species, the tundra may lose specialists under increasing climate stress while being progressively invaded by boreal taxa from the south.

Beyond the scope of our model, tundra plants might integrate into the invading boreal forest if they share ecological functions with boreal species that have limited dispersal and have not yet expanded. Such replacement has been observed among congeners (Schmidt et al., 2012). However, competition from immigrating boreal species could accelerate tundra decline (Callaghan et al., 2004), and competitive exclusion may drive extinctions even more strongly than temperature increase (Pykälä, 2017). Over time, vegetation in boundary regions may increasingly resemble boreal communities, leading to a shrinking tundra area.

Spatial configuration also shapes range shifts. In regions such as Siberia, tundra cannot expand northward because no land lies beyond (Pearson et al., 2013). Proximity to the sea across much of the tundra further restricts expansion, contributing to its decline even in the unconstrained scenario. Our model predicted greater northward migration of the Nearctic boreal forest compared to the Palearctic, reflecting the longer northward extent of the Nearctic landmass. In contrast, Palearctic species colonized more cells overall because the Palearctic landmass is larger, so more area becomes suitable at the same latitude. Thus, in addition to dispersal constraints, geographic constraints play a major role in shaping biome-scale range shifts.

### 4.4 Further implications

The predicted vegetation shifts are likely to occur more slowly and less strongly in reality, as our models did not account for extirpation nor establishment lags. Much of the change we observed results from predicted extirpations under declining climatic suitability; assuming no extirpations led to much weaker compositional change. In addition, colonization is not determined by dispersal alone but can be slowed by processes such as competition creating establishment lags.

For future studies, we recommend incorporating dispersal constraints, which were shown to substantially impact species distributions. Additionally, extirpation and establishment lag times, or, in case of mechanistic models, species interactions and species-specific time until maturity that can cause these time lags, should be included as these factors also impact the realized niche of species. To predict changes of the entire ecosystem, integrating animals would be necessary, as herbivory could counteract the predicted dynamics of plant distributions.

As climatic stress and competition of invasive species but also other factors like land use change and pollution are threatening biodiversity in the Arctic (Ksenofontov et al. 2018), innovative conservation measures are crucial. For instance, the protection of climate refugia, areas that remain relatively stable despite broader environmental change, is important, as these refugia offer vulnerable species the chance to persist and potentially re-expand (Keppel et al. 2012). Moreover, a proactive strategy to protect threatened species is assisted migration (aka assisted colonization, managed translocation), the deliberate relocation of species to areas where they are more likely to survive under future climate conditions (Hällfors et al. 2014). Although this approach is controversially discussed, it might be reasonable to bring the seeds of poor dispersing tundra species further north, where plants are barely established today, as the tundra is threatened to lose area at its southern edge which becomes invaded by boreal forest species. However, careful consideration of ecological impacts and ethical implications is essential when implementing such interventions (cf. Hunter 2007; Richardson et al. 2009; Minteer and Collins 2010; Aubin et al. 2011; Hewitt et al. 2011; Sáenz-Romero 2021).

### 4.5 Conclusion

By comparing a dispersal-unconstrained scenario with one incorporating species-specific dispersal limits, we show that dispersal ability strongly shapes future species distributions and the capacity to track shifting climate niches. While new habitats are projected to emerge mainly at higher latitudes, higher elevations may serve as more accessible refugia. Despite potential niche gains, overall Arctic plant richness is expected to decline under climate warming because many species cannot reach suitable areas. Together, colonization of new sites and loss of old ones are predicted to drive major shifts in community composition and to transform the boreal forest–tundra boundary.

## Supporting information

Appendix overview

ODMAP protocol

summary current climate

climate summary

SDM parameters

traits and dispersal rates

Validation

How to excess the SDM output

