## Appendix overview for "Different dispersal rates and declining climate suitability shape future vegetation compositions across the Arctic"

**Table of contents - Appendix**

1. **S1 ODMAP**	2
2. **S2: Further model information**	2
3. **S3: Species parameters data**	6
4. **S4: Trait and dispersal data**	6
5. **S5: SDM Output**	6
6. **S6: Validation of dispersal rate**	7
7. **S7: Validation of species distribution maps**	8
8. **S8: Validation Ulukhaktok	11**
9. **S9: Cell classification	12**
10. **S10: Boreal indicator tree species list	13**
11. **S11: Beta diversity	13**
12. **S12: Other scenarios	14**
13. **References**	22

**S1 ODMAP protocol see ODMAP.csv** (see for detailed model description)

**S2: Further model information**

**Maxent settings**

To run the dismo::maxent() function, we used default settings for the arguments (args = c('responsecurves=FALSE', 'jackknife=FALSE', 'pictures=FALSE', 'autofeature=FALSE', 'linear=TRUE', 'quadratic=TRUE', 'product=TRUE', 'threshold=TRUE', 'hinge=TRUE', 'betamultiplier=1'). Maxent uses all features (such as linear, quadratic, product) for occurrences > 80, as in our case, extracting the most useful ones (Merow et al. 2013).

**Binary threshold**

There are several thresholds to transfer the continuous output (0 - 1) into binary values (0 or 1) by selecting a value in the receiver operating characteristic (ROC) curve above which the specific species is considered present and below it is considered absent (Phillips et al., 2006). Merow et al. (2013) recommend neither arbitrary thresholds (such as 0.5, as choosing values sensefully can depend on density or prevalence of the population), nor specificity-based measures (as they are calculated based on truly predicted absences which does not work for background data). Sampling bias is considered the largest challenge of presence-only models (Merow et al., 2013), so we decided to use the prevalence threshold “10 percentile training presence” which describes that 90 % of all input occurrence points are included in the probabilities above the threshold. The value of this threshold was in the middle of the range of nine MaxEnt thresholds and represented the present distributions of common species well, opposed to higher thresholds which left out important areas of their occurrences.

**Bioclim variables**

https://www.worldclim.org/data/bioclim.html, 28/01/2025

BIO1 = Annual Mean Temperature

BIO2 = Mean Diurnal Range (Mean of monthly (max temp - min temp))

BIO3 = Isothermality (BIO2/BIO7) (×100)

BIO4 = Temperature Seasonality (standard deviation ×100)

BIO5 = Max Temperature of Warmest Month

BIO6 = Min Temperature of Coldest Month

BIO7 = Temperature Annual Range (BIO5-BIO6)

BIO8 = Mean Temperature of Wettest Quarter

BIO9 = Mean Temperature of Driest Quarter

BIO10 = Mean Temperature of Warmest Quarter

BIO11 = Mean Temperature of Coldest Quarter

BIO12 = Annual Precipitation

BIO13 = Precipitation of Wettest Month

BIO14 = Precipitation of Driest Month

BIO15 = Precipitation Seasonality (Coefficient of Variation)

BIO16 = Precipitation of Wettest Quarter

BIO17 = Precipitation of Driest Quarter

BIO18 = Precipitation of Warmest Quarter

BIO19 = Precipitation of Coldest Quarter

**Importance of predictor variables**

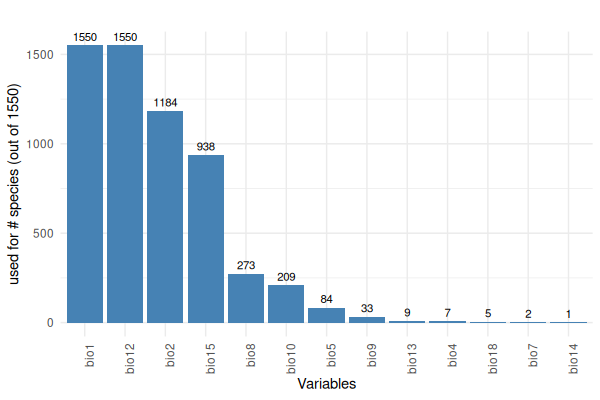

***Figure S2.1: For each environmental variable, the number of species distribution models in which it has been included is given. Variables on the right are more important as they have been used to set up the model of all 1550 species (Note that we included bio1 (annual mean temperature) and bio12 (annual precipitation) by default as basic variables for all species and added variables which were not collinear).***

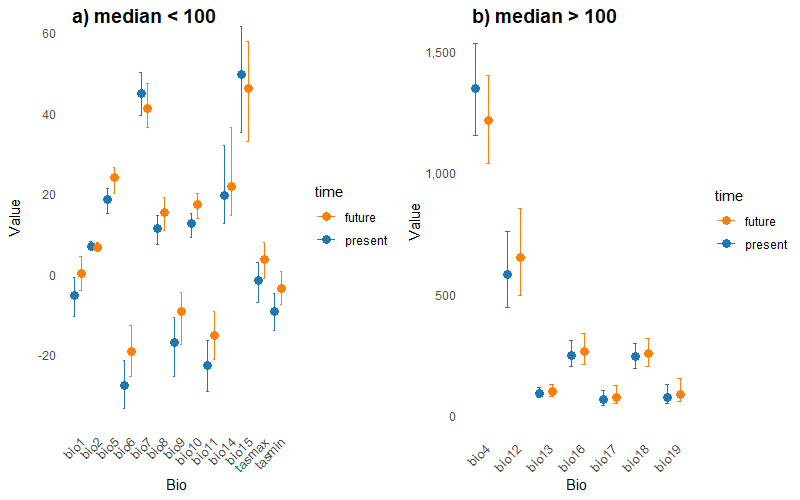

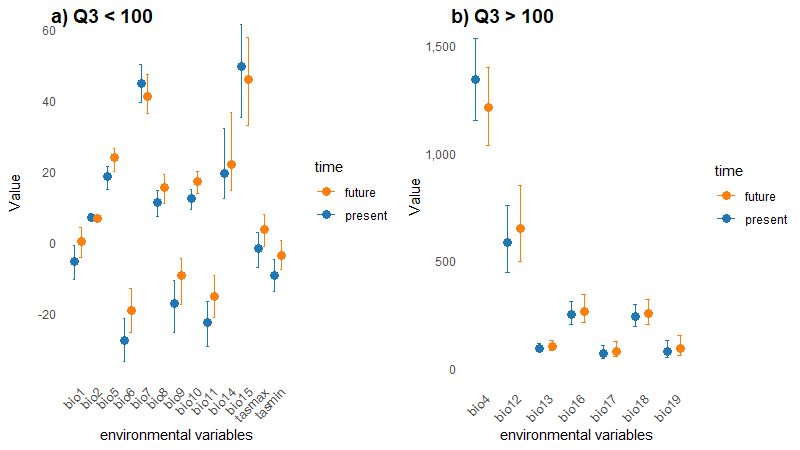

***Figure S2.2: Median (dots) and 3rd quantile (75 % percentile) (segments) of environmental variables for present (blue) and future (2100, scenario ssp585) (orange). For better visualization of the values, it is splitted in two plots, for values of the 3rd quantile below and above 100. You can confer the meaning of bio1-bio19 from the overview table above the plot.***

**Spatial filtering**

In addition to grid-based thinning, we performed spatial filtering on our data to maximize equal spatial representation, as occurrence points were skewed towards Europe. We here show why we included this additional step by comparing the evaluation metrics of the “bias_correction” method (with spatial filtering step, as used in the paper) to the “gloabal_bias” method (without spatial filtering step, not used in the paper).The boxplots are arranged by descending median of the global_bias results. It can be seen that the global_bias metrics decrease from high values for the well-studied regions Sweden, Finland and Norway towards much lower values for regions with less sampling records (e.g. Mongolia). The bias_correction results show a smaller variation of the median over the regions; the smallest AUCs of all regions are higher than the smallest AUCs for the global_bias results. For the regions with less sampling records the bias_correction results show higher metrics than the global_bias results. For the well-studied areas Sweden, Finland and Norway the global_bias results are better than the bias_correction results. For all regions evaluated together, the metrics are higher for the global_bias results, but this is probably due to the higher performance in well-studied regions. As we aim for a large-scale prediction of the entire circumpolar area, we wanted to mitigate the effects of sampling bias aiming to meliorate predictions in regions with less sampling records. We thus received better model performance over all regions when applying the bias_correction method and decided to use this method for the result presented in our paper.

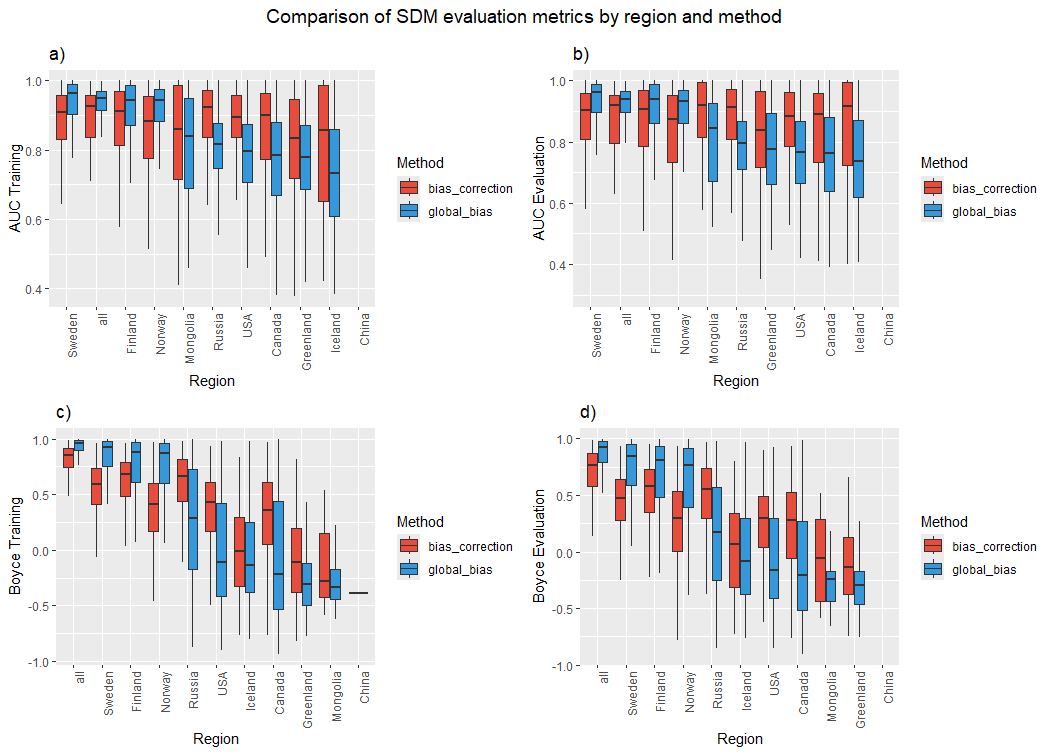

***Figure S2.3: Comparison of SDM evaluation metrics by region and method. a) AUC based on training data, b) AUC based on evaluation data, c) Boyce index based on training data and d) Boyce index based on evaluation data. The method indicates the decision to stay with one step of (local) spatial filtering on the 5 km x 5 km environmental grid (global_bias) or to include another step of sampling bias correction by applying a second step of (large-scale) spatial filtering in the model code (bias_correction) to avoid overfitting on Sweden, Finnland and Norway.***

**S2.4_summary_table_env_curr.csv**
Overview of environmental variables at present.

**S2.5_climate_summary.csv**
 Overview of environmental variables in 2100, scenario ssp585.

**S3: SDM parameters**

For species list, binary threshold, number of occurrences and background points, AUC, Boyce Index and included predictor variables for each species see appendix table **S3_SDM_parameters.csv.**

**S4: Trait and dispersal data**

For the collected traits and the assigned dispersal distance class for each species see appendix table
**S4_traits_dispersal_rates.csv**

**S5: SDM Output**

For each species, present distribution maps (1980 – 2010), occurrence points and a table with evaluation metrics are compiled in the html file
**S5_SDM_output.html**

**S6: Validation of dispersal rate**

***Table S6.1: Dispersal rates [m/year] for different arctic plant taxa estimated in our study following Lososová et al. (2023) compared to dispersal rates from literature given with source and additional details.***

| **taxa** | **Our estimate  [m/year]** | **Literature value [m/year]** | **source** | **details** |
| --- | --- | --- | --- | --- |
| trees | 150 - 1500 | 10^2^ – 10^3^ | (Clark et al., 1998) | end of the Pleistocene |
| trees | 1500 maximum | 1600 | (Cunze et al., 2013) | average of their 140 European plant species with different dispersal modes |
| all 1174 species average | 336 | 270–380 | (Snell, 2014) | past rates that  were assumed for the whole landscape |
| *Acer spp.* | 150 | 141 | (Snell, 2014) |  |
| *Picea spp.* | 150 | 140 | (Clark et al., 1998) | meta study |
| *Galium aparine* | 5 | >2 | (Matlack, 1994) |  |
| *Asarum europaeum (our)/ canadense (lit.)* | 15 | 12.5 | (Cain et al., 1998) | 200 km in 16,000 years |

The comparison to literature shows that the dispersal rates we assigned following Lososová et al. (2023) are appropriate estimates concerning single species as well as regarding the overall magnitude.

**S7: Validation of species distribution maps**

Unfortunately, for most species there are no large-scale distribution maps for arctic vegetation available. Best data source for that is GBIF.org, however we already used the available data on GBIF.org for calibration. We compared distribution maps from studies for single species for certain regions. Here are some examples:

*Picea glauca*

Ohse et al., 2009

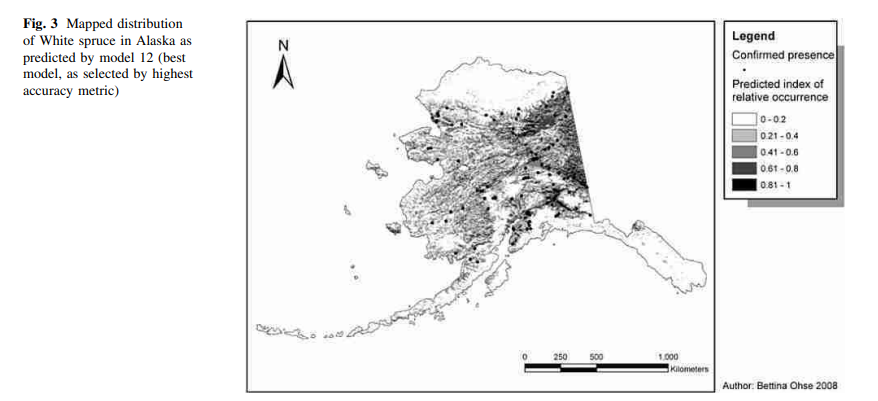

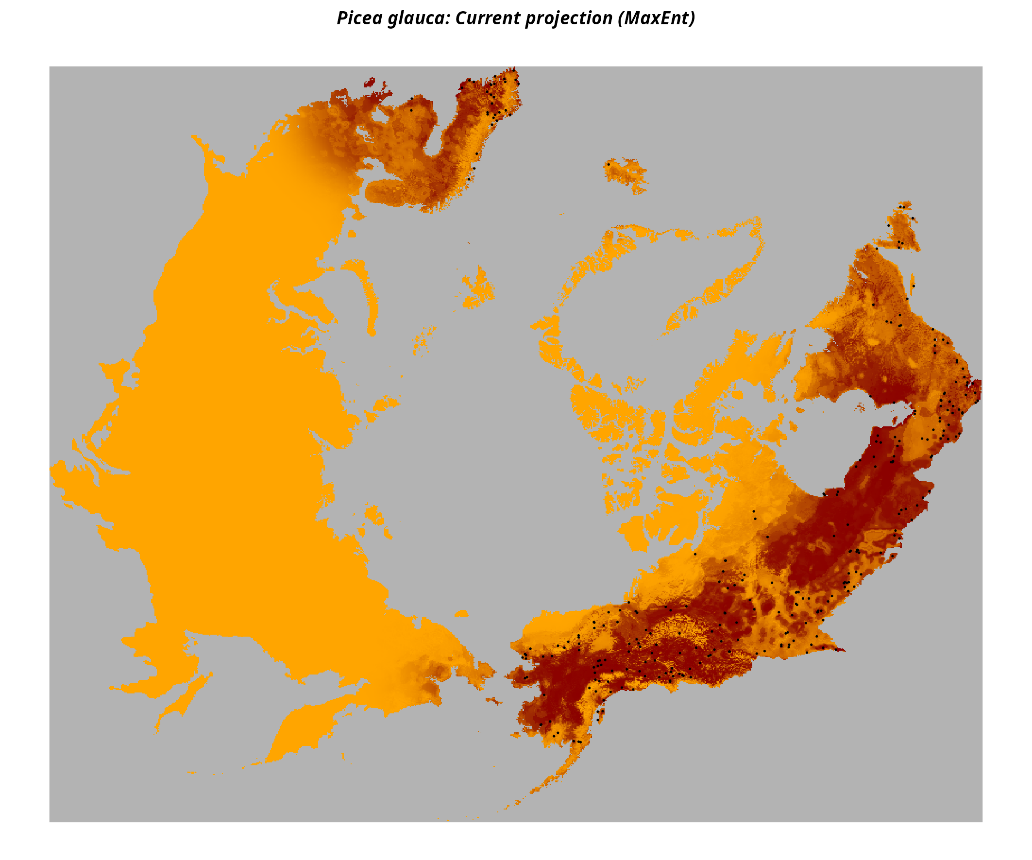

***Figure S7.1: Comparison of literature distribution (above) with our species maps (below) for Picea glauca.***

*Larix (gmelinii)*

Kruse & Herzschuh, 2022

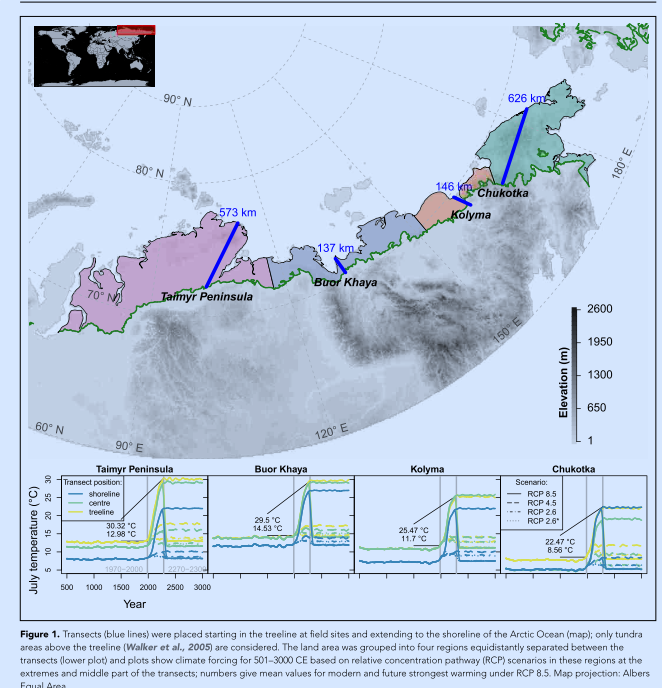

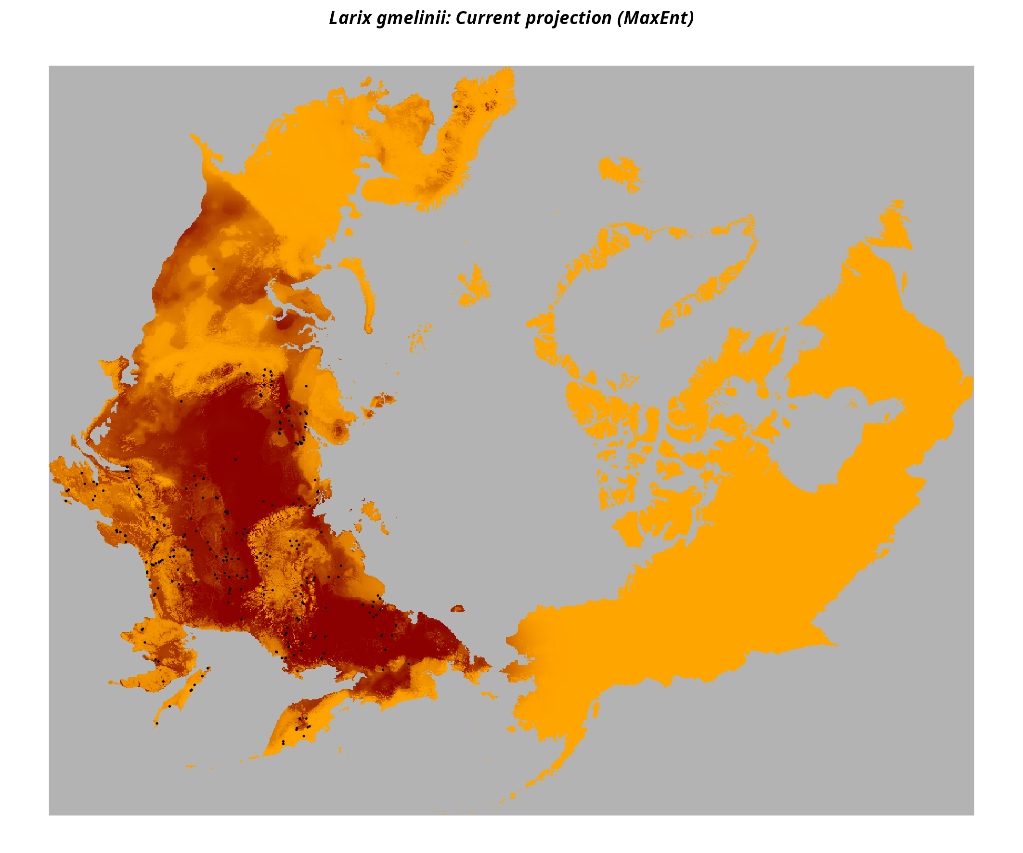

***Figure S7.2: Comparison of Larix spp. literature distribution (above) with our species maps (below) for Larix gmelinii.***

*Eriophorum vaginatum*

Curasi et al., 2022

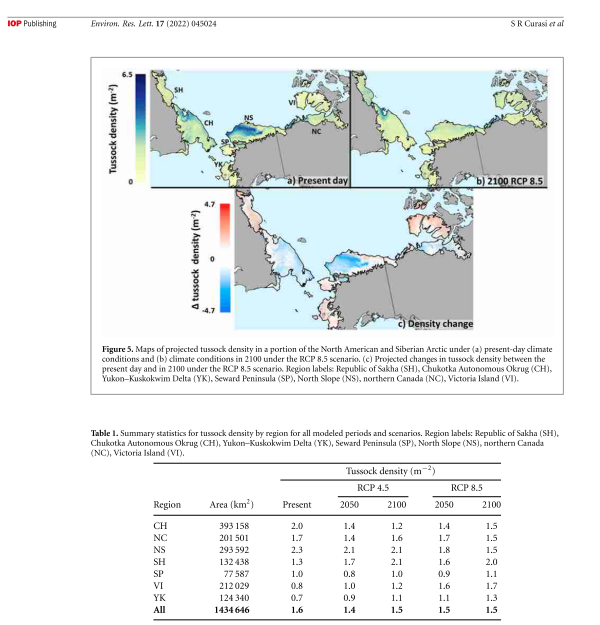

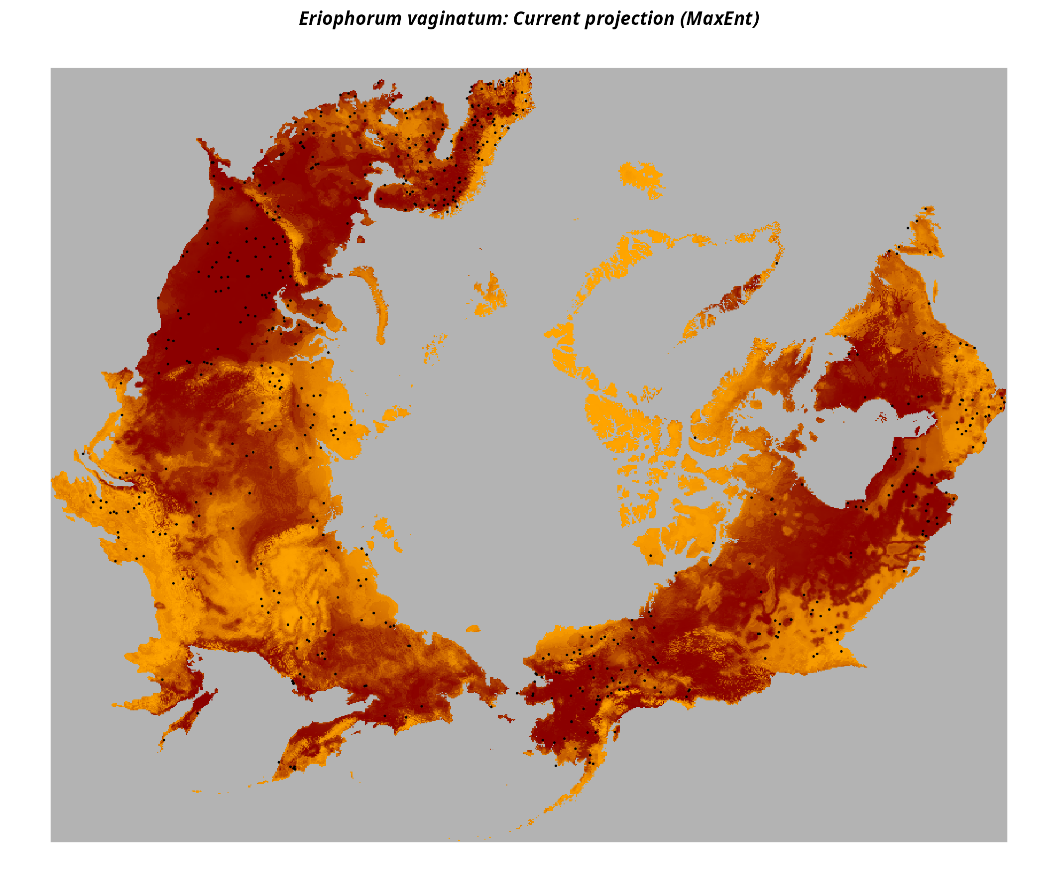

***Figure S7.3: Comparison of literature distribution (above) with our species maps (below) for Eriophorum vaginatum.***

**S8: Validation Ulukhaktok**

Aiming for a validation in the field, during an expedition to Ulukhaktok (Northwestern Territories, Canada) in summer 2025 (Figure S11.1, left), plant species were documented by vegetation mapping and collection of specimens for an herbarium record and additionally observed during hiking. Our species list for this area included 49 different plant species out of which 36 species were also included in this study. For these species we extracted the suitability and binary result (presence/absence) for the time slice 2010 – 2040 and all three scenarios (ssp126, ssp370, ssp585) at the coordinates of six sites we visited (for coordinates see Table S11.1; for extracted data see the csv-file S8.2_ Ulukhaktok_validation_all3scen_sites.csv). Comparing our field species list for Ulukhaktok to the species that were predicted present by the model for at least one of these sites, 32 species were predicted correctly, while five of the recorded species were predicted as absent at all sites. Our model hence showed an agreement of 86% with the field records from Ulukhaktok.

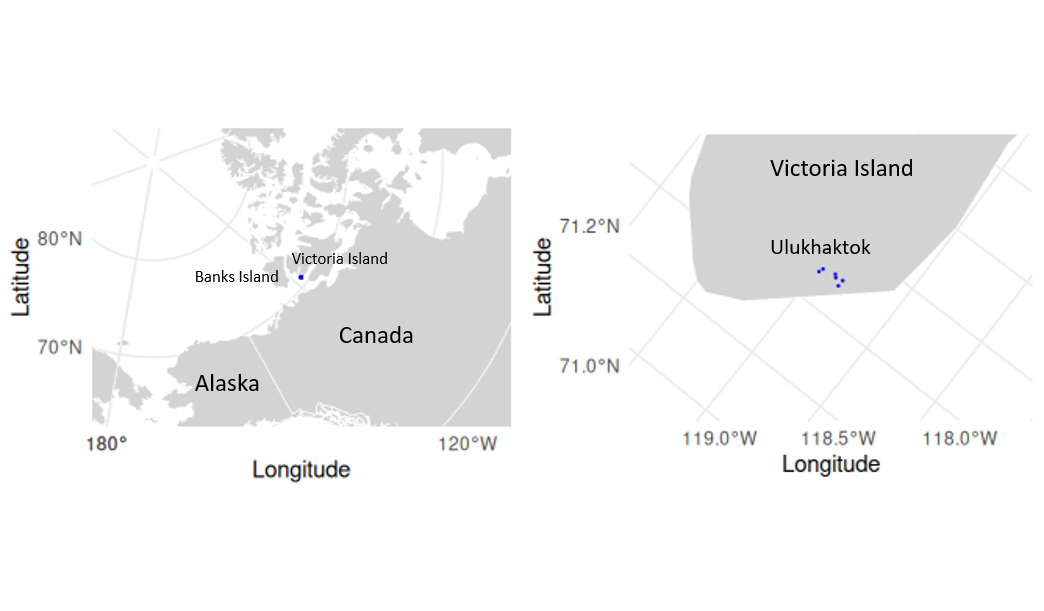
**Figure S8.1: Locations of sampling sites in Ulukhaktok (Victoria Island, Northwestern Territories, Canada) on large (right) and small (left) scale.**

**Table S8.1: Coordinates of sites visited during vegetation mapping and collection of specimens for an herbarium record in summer 2025 in Ulukhaktok (Northwestern Territories, Canada).**

| Longitude | Latitude | site |
| --- | --- | --- |
| -117.7537 | 70.7542 | 217 |
| -117.7378 | 70.7599 | 218 |
| -117.7606 | 70.7895 | 220 |
| -117.7911 | 70.7932 | 219 |
| -117.7874 | 70.7382 | 221 |
| -117.7412 | 70.7375 | 222 |

**S8.2_ Ulukhaktok_validation_all3scen_sites.csv**

Suitability and binary result (presence/absence) for the time slice 2010 – 2040 and all three scenarios (ssp126, ssp370, ssp585) at the coordinates of six sites we visited

**S9: Cluster classification**

We matched our cell clusters (vegetation communities) with the respective WWF ecoregions (Olson et al., 2001) (which we also used during the model setup to download GBIF data for that region).

WWF_REALM2 == "Nearctic" or "Palearctic" (realms)

WWF_MHTNAM == "Boreal Forests/Taiga" or "Tundra" (biomes)

Agreement of our classification:
percentage of correctly classified cells per cells in the WWF ecoregion

Nearctic + Boreal Forests/Taiga: 89% (wrong tundra,11%)
Palearctic + Boreal Forests/Taiga: 93% (wrong tundra, 7%)
Tundra: 63 % (wrong: Palearctic boreal forest 30%, Nearctic boreal forest 7%)
**Note than we do not claim our “no tree” area to be the tundra area, but rather a part of it,
as it only covers 63 % of the tundra area.**

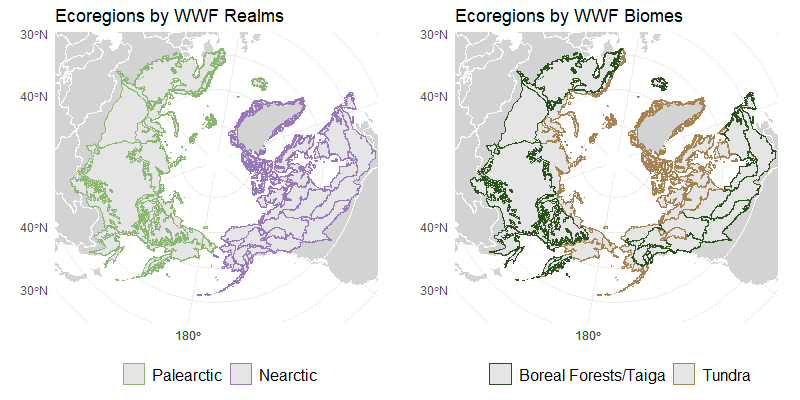

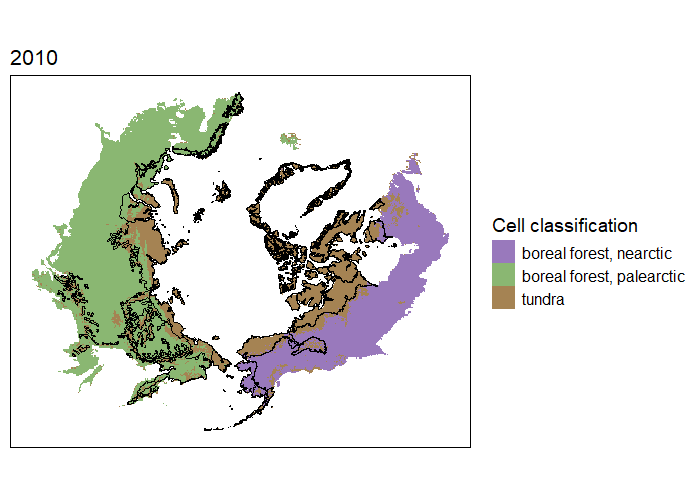

***Figure S9.1: Biomes and realms in the Arctic according to the ecoregions*** ***dataset (Olson et al., 2001) (above). Main cell clusters in our study named following the ecoregion classification, showing tundra boundaries according to Olson et al. (2001) in black lines (below).***

**S10: Boreal indicator tree species list**

**S10_boreal_tree_list.csv**

Overview over all tree species and the ones selected as typical boreal tree species to model the distribution of the boreal forest

**S11: Beta diversity**

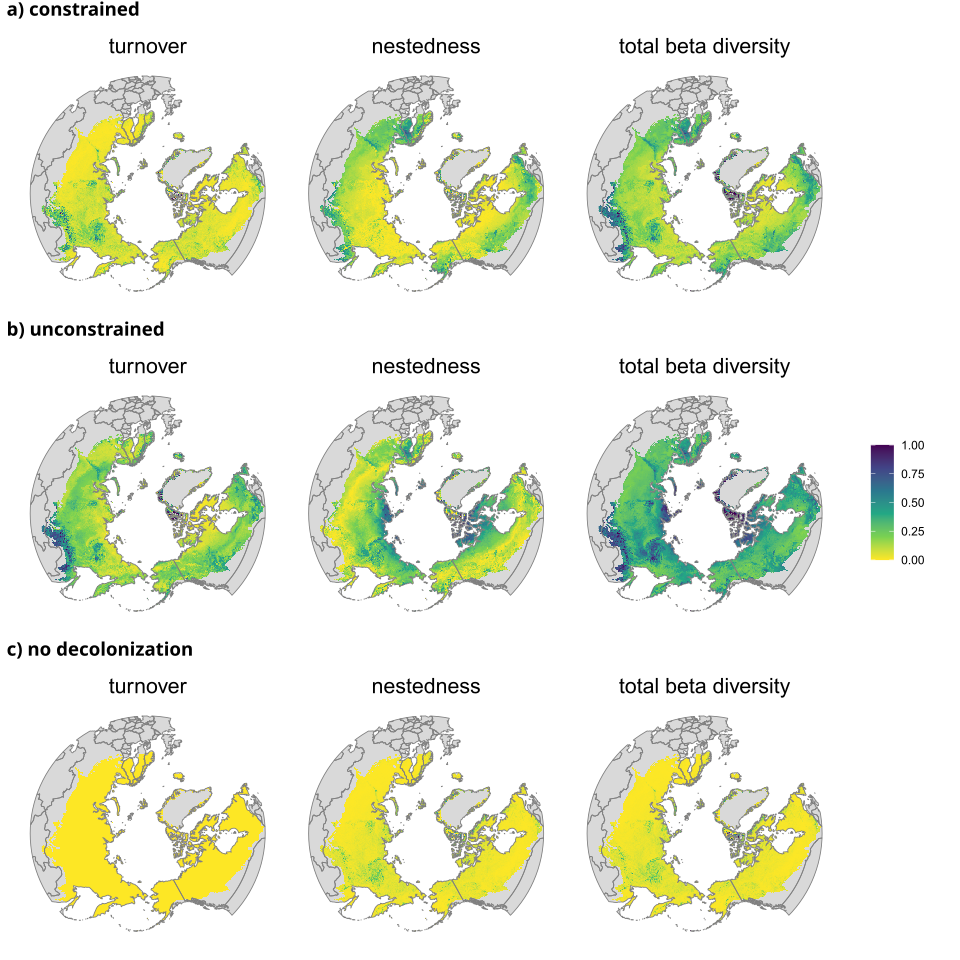

***Figure S11.1:*** ***Contribution of turnover (proportion of beta diversity that results from species replacement between sites) and nestedness (proportion of beta diversity that results from species loss or gain without replacement) to beta diversity and total beta diversity (estimated with package ‘betapart’ (v.1.6.1), Baselga et al. (2025)) are shown for a) dispersal constrained, b) unconstrained and c) no decolonization scenario.***

**S12: Climate scenarios ssp126 and ssp370**

ssp126	ssp370

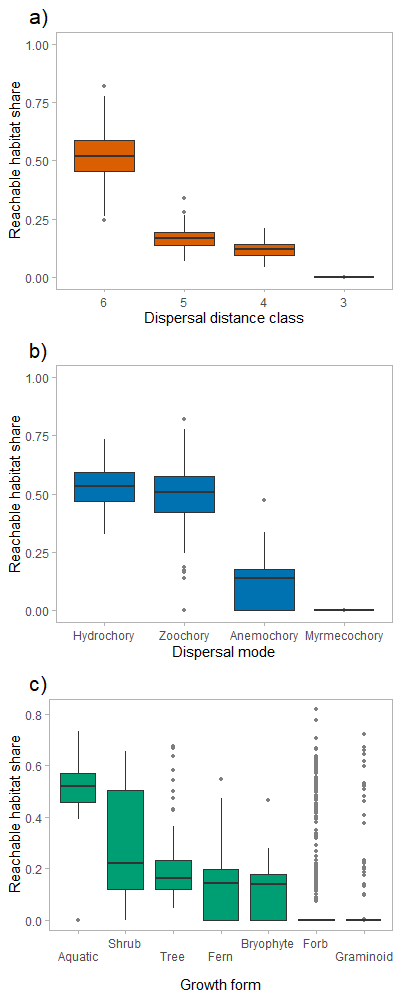

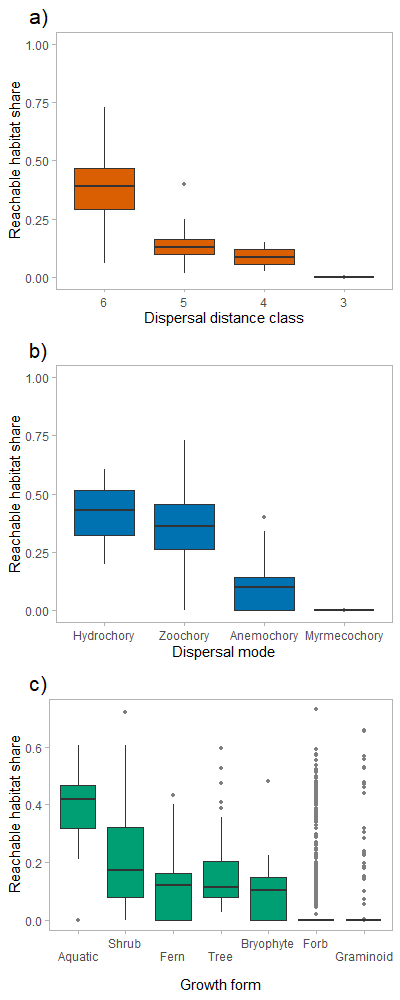

***Figure 3:*** *Reachable habitat share, i.e. new colonized cells in 2100 (climate scenario ssp585) compared to 2010 for the dispersal constrained scenario divided by the new colonized cells in 2100 (climate scenario ssp585) compared to 2010 for the unconstrained scenario for the (a) dispersal distance class, (b) dispersal mode and (c) growth form.* *Climate scenario ssp126 (above) and ssp370 (below).*

ssp126

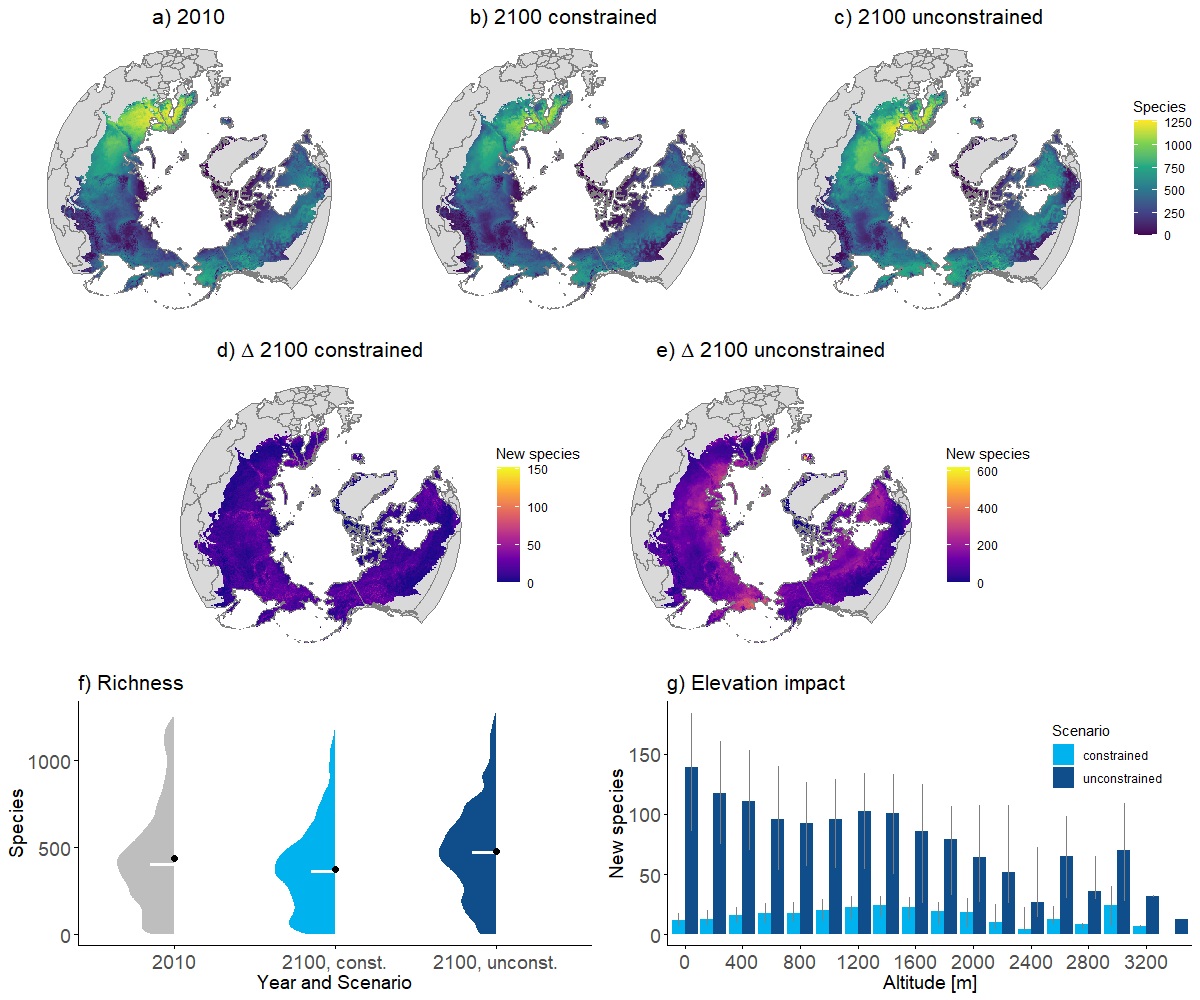

ssp370

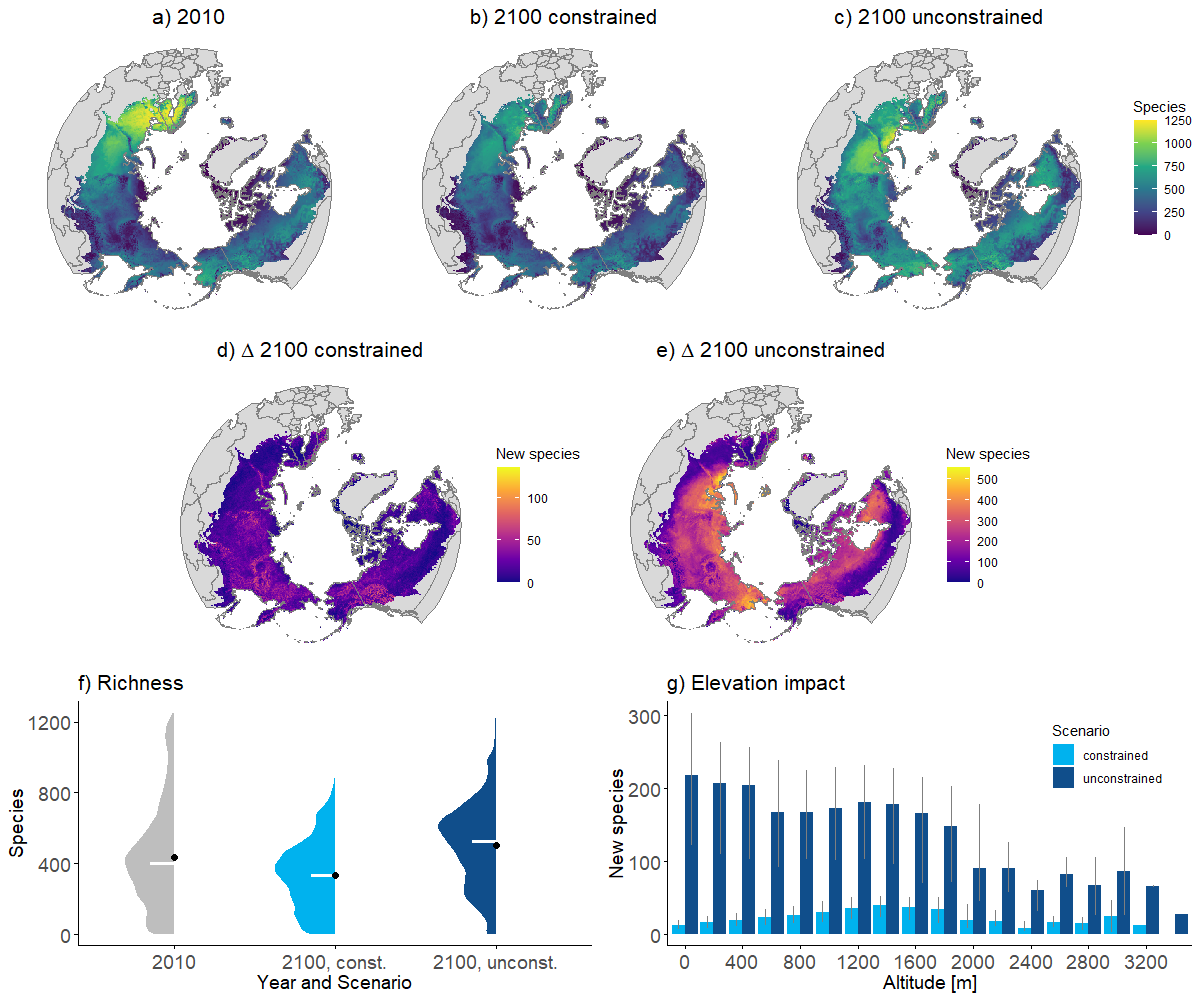

***Figure 4:*** *Number of all studied 1550 arctic plant species per cell for (a) present and for 2100 (last of three time slices) for climate scenario ssp585 (maximum scenario out of three) for the (b) species-specific dispersal constrained and (c) unconstrained scenario. Number of only new colonizing species by 2100 (climate scenario ssp585) compared to 2010 for the (d) dispersal constrained and (e) unconstrained scenario. The plots show the sum of the distribution maps of all 1550 species (a - c) and the difference of the sum between 2100 and 2010 (d - e). All maps are presented in the Lambert Azimuthal Equal Area projection. (f) Number of all species per cell as half violin plot for the data shown in (a), (b) and (c). (g) shows the number of new species over different altitudes for the constrained and unconstrained scenario. The grey error bars show the 25% and 75% quantiles. Climate scenario ssp126 (above) and ssp370 (below).*

ssp126

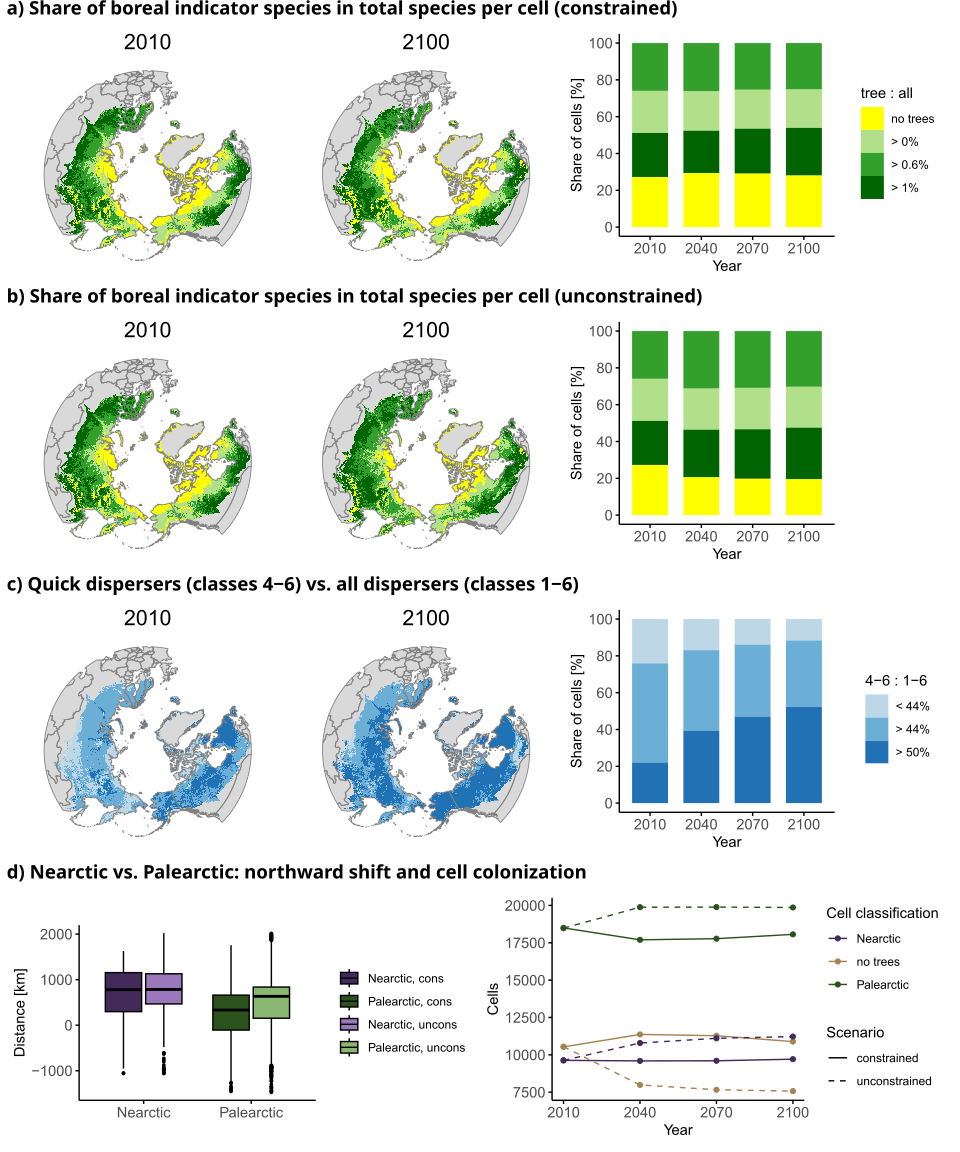

ssp370

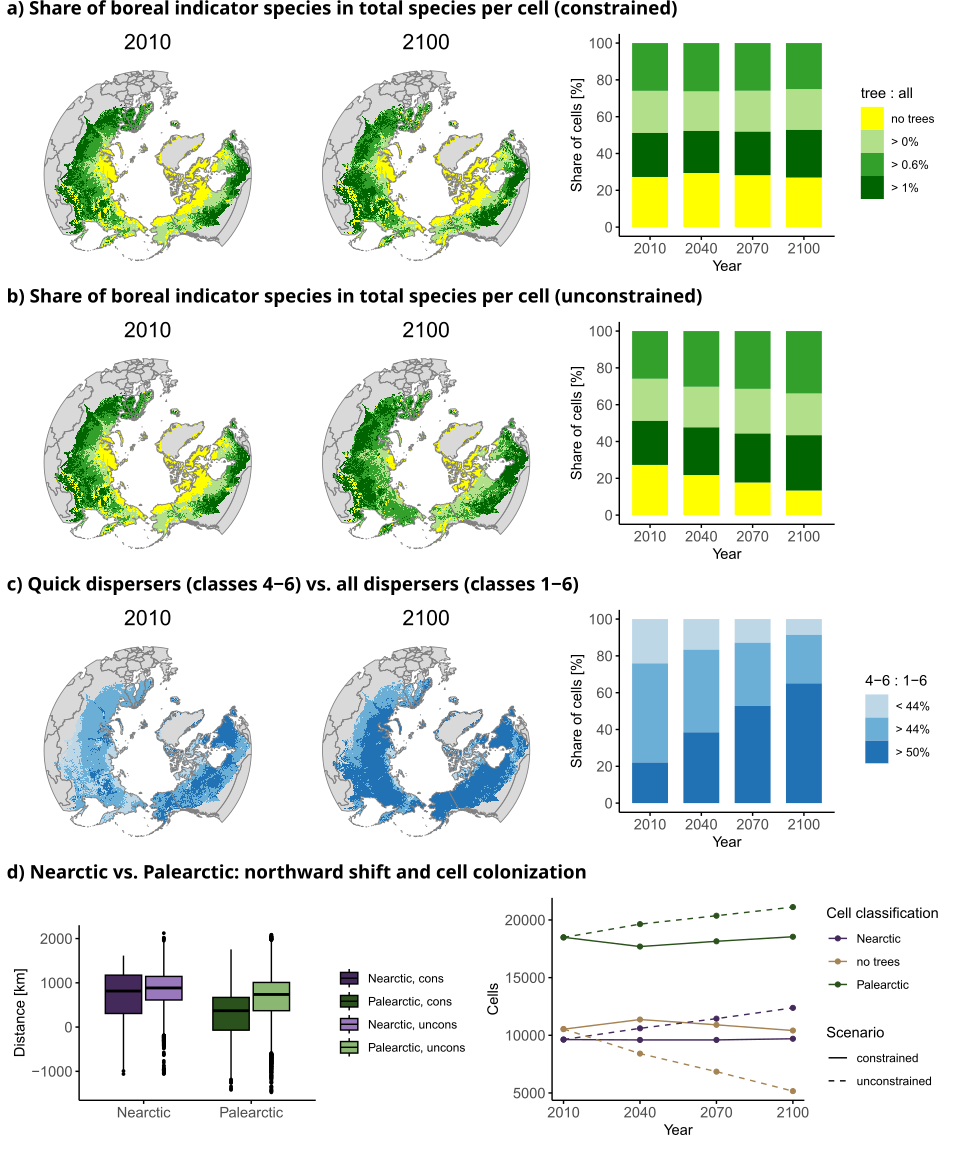

***Figure 5****: Shift of boreal forest - tundra boundary over time (2010, 2040, 2070, 2100) for climate scenario ssp585 (maximum of three scenarios) visualized as share of boreal indicator tree species in total number of species per cell for (a) dispersal constrained scenario and (b) unconstrained scenario.c) Share of quick dispersing species (belonging to dispersal distance classes 4-6) in all species per cell (dispersal distance classes 1-6)* *(d) Northwards shift of the boreal forest regions in Nearctic and Palearctic from 2010 to 2100 for constrained and unconstrained scenario (left) and number of cells occupied by the cell conglomerates over time for the dispersal constrained scenario (solid line) and unconstrained scenario (dashed line). The cell conglomerates were named according to their geographic position in the realms and the similarity to the biomes of the boreal forests (incl. taiga) and the Arctic tundra (“no trees” (Olson et al. 2001), were the presence of at least one tree species defines the “boreal forest” hence the underlying data is the binary version of the ratio shown in (a) and (b). Climate scenario ssp126 (above) and ssp370 (below).*

ssp126

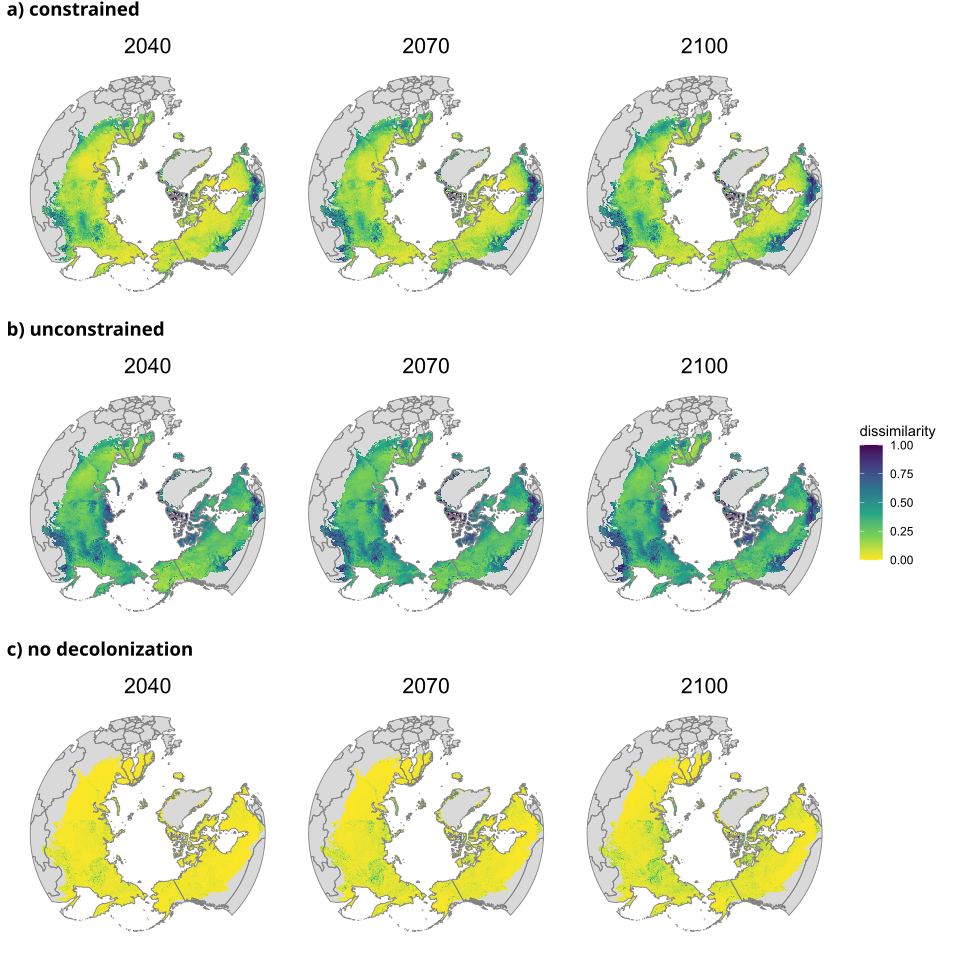

ssp370

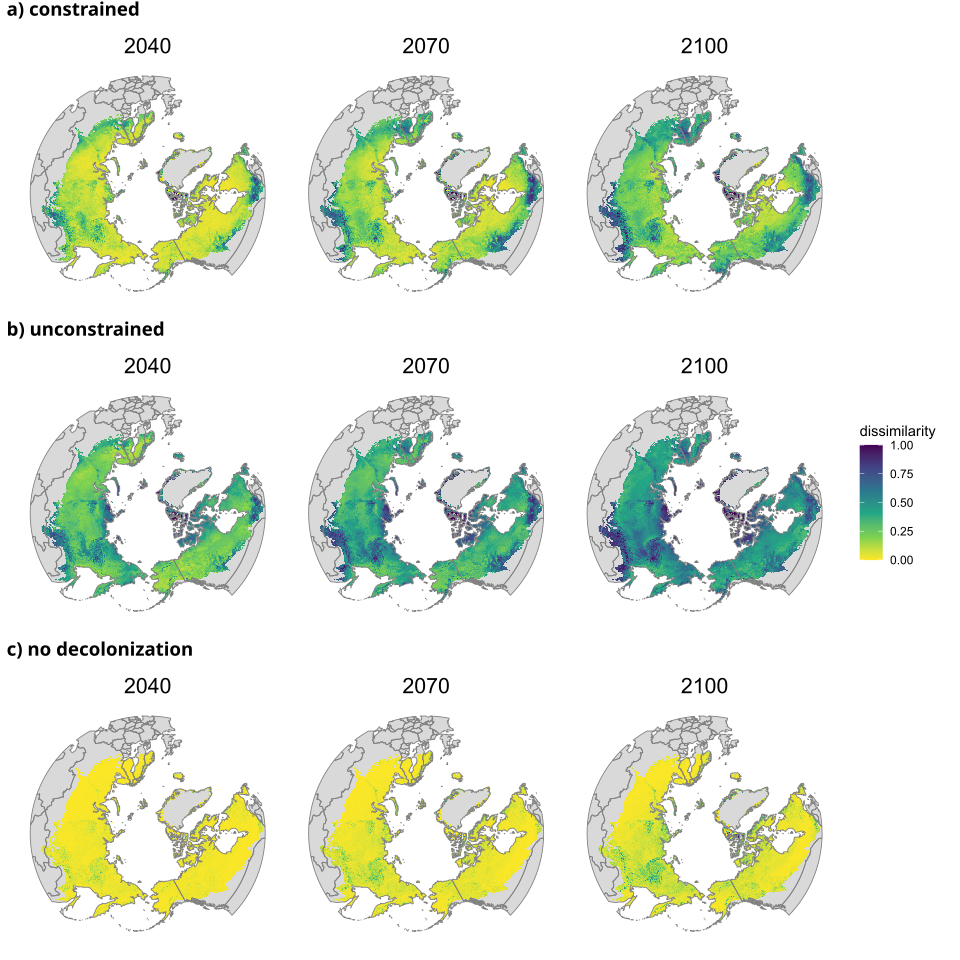

***Figure 6:* *Impact of dispersal vs. extirpation: Dissimilarity (Jaccard index) of species composition compared to 2010 over time for (a) constrained, (b) unconstrained and (c) no decolonization scenario (based on constrained scenario).*** *Here, no decolonization means that species may only colonize new cells but may not disappear if the climate niche becomes unsuitable (see discussion for potential reasons and implications).* Th*e Jaccard dissimilarity index ranges between 0 “total similarity” and 1 “total dissimilarity”. Climate scenario ssp126 (above) and ssp370 (below).*
