## Supplementary material for "Different dispersal rates and declining climate suitability shape future vegetation compositions across the Arctic": How to excess the SDM output

The S5_SDM_output.html is 630 MB and was therefore too large for upload.
You are welcome to contact us in case you are interested, and we will send it to you.
Please note that it will shortly be published along with the Paper in the Journal of Biogeography.
